# Structural and functional examination of *Neisseria gonorrhoeae* TfpC reveals a dynamic two-domain architecture required for type IV pilus function

**DOI:** 10.64898/2026.11.06.764878

**Authors:** Kathleen R. Nicholson, Maria Baczynska, Eliza Płoskoń, Peter Simpson, Alain Origioni, Christian D. Lorenz, H Steven Seifert, James A. Garnett

## Abstract

The *Neisseria gonorrhoeae* (Gc) Type IV pilus (T4p) is a critical apparatus for colonization and virulence. This molecular machine resides at the host-pathogen interface and mediates adherence to host cells, twitching motility, natural transformation, and resistance to oxidative agents. Previously, we identified a DUF4124 family, T4p component, TfpC. TfpC is required for full T4p elaboration, and deleting the *tfpC* gene reduced natural transformation competence and resulted in an under-piliated colony morphology. Our earlier models suggested TfpC is composed of an N-terminal transmembrane helix and a proline-rich disordered region followed by a C-terminal coiled-coil. Here, we used targeted structural and genetic approaches to define TfpC’s molecular structure. Our new data show that TfpC’s N-terminal domain forms a novel, dynamic three-stranded β-sheet structure. These data indicate that the flexible disordered region acts as a linker between the N-and C-terminal domains. Functionally, we illustrate TfpC requires an acidic face within the C-terminal coiled-coil domain to mediate pilus biogenesis and function. This study characterizes a unique protein fold and family and defines the contributions of TfpC’s domains to its function.

## Introduction

*Neisseria gonorrhoeae* (Gc) is a Gram-negative bacterium responsible for most gonorrhea infections in humans (*WHO Guidelines for the Treatment of Neisseria Gonorrhoeae*, 2016). Gc Type IV pili (T4p) are a major colonization virulence factor responsible for adherence to host cells, which initiates colonization of the host (Merz & So, 2000; Sparling, 1966). T4p are common molecular machines in almost all Gram-negative bacteria and a subset of Gram-positive bacteria (Craig & Li, 2008). These structures support various elements of bacterial physiology, including natural transformation, twitching motility, resistance to oxidative agents, and perhaps most importantly, virulence (Merz & So, 2000; Sparling, 1966; Stohl et al., 2012, 2013).

T4p are assembled by a multiprotein machine that spans the cell envelope (Chang et al., 2016; Craig et al., 2019). The major pilin PilE is inserted into the inner membrane by the Sec machinery, processed by the prepilin peptidase PilD, and then mature pilins are retained in the membrane through their conserved hydrophobic N-terminal α-helix (Craig et al., 2019; Obergfell & Seifert, 2016). The extension ATPase PilF facilitates the movement of these subunits to the base of the growing fiber at the inner membrane platform protein PilG, which are then extracted and incorporated into the helical filament (Chang et al., 2016; Kolappan et al., 2016). The retraction ATPases PilT and PilU depolymerize the fiber, and cycles of extension and retraction generate the forces required for twitching motility and DNA uptake (Merz et al., 2000). The fiber crosses the periplasm and outer membrane through the PilQ secretin, which is aligned with the inner membrane components by a subcomplex of PilM, PilN, PilO, and PilP (Berry et al., 2012; Georgiadou et al., 2012). The minor pilins ComP, PilV, and PilX are incorporated into the fiber and contribute to priming of assembly, aggregation, and natural transformation (Cehovin et al., 2013; Helaine et al., 2007; Wolfgang et al., 1999). Accessory proteins interact with this core machinery, with PilW required for PilQ oligomerization and fiber stability (Carbonnelle et al., 2005; Szeto et al., 2011), and the peptidoglycan-binding protein TsaP required for surface assembly of pili (Siewering et al., 2014). While these core components are well defined, several other proteins that are required for full piliation have no assigned position within the machine.

We and others previously published the identification of a novel T4p component of *N. gonorrhoeae* and *Neisseria meningitidis,* respectively, and we named this component TfpC (Hu et al., 2020; Muir et al., 2020). TfpC is required for full Gc piliation, and strains lacking TfpC have fewer and shorter pili and reduced transformation competence, indicating TfpC is important for full function of the T4p (Hu et al., 2020). The N-terminus of TfpC is annotated as a Domain of Unknown Function (DUF4124), while the C-terminus is a predicted coiled-coil domain, and these domains are connected by a long proline-rich unstructured region. No experimentally determined structure has been reported for any DUF4123-domain protein. Our previously published TfpC structural model suggested the N-terminus forms a transmembrane helix that may insert into the inner membrane (Hu et al., 2020). This would localize TfpC in the inner membrane and constrain the position of its C-terminal coiled-coil, whereas a soluble periplasmic protein could engage separate partners at either end of its disordered linker. Despite the importance of TfpC for T4p expression and function, its localization within the cell envelope has been inferred from sequence-based prediction alone, and no binding partner has been identified (Hu et al., 2020; Muir et al., 2020). In this study, we refined the structure and dynamics of TfpC and used genetic approaches to perform structure-function analyses showing both domains are important for pilus function and highlighting potential binding interfaces.

## Results

### Overall structure of the TfpC N-terminal region

Our previous structural model of full-length TfpC was derived from co-evolution analysis and predicted the N-terminus to be a single helix (Hu et al., 2020). However, the Jpred4 secondary structure server (Drozdetskiy et al., 2015) instead predicted the N-terminus to be composed of three β-strands (residues 1–27 of the mature protein) (**Supplementary Figure 1,2**). To resolve this discrepancy, we carried out solution-state Nuclear Magnetic Resonance (NMR) spectroscopy on TfpC, beginning with a peptide spanning residues 1–27 of its N-terminal region (TfpC_NTD27_) (**Supplementary Figure 1**). Analysis of the resulting ^1^H-^1^H Total Correlation Spectroscopy (TOCSY) and ^1^H-^1^H Nuclear Overhauser Effect Spectroscopy (NOESY) spectra at 25°C revealed limited chemical shift dispersion and very few medium-or long-range Nuclear Overhauser Effect (NOE) correlations, indicating that the isolated peptide is predominantly disordered in solution (**Figure 1A, Supplementary Figure 3**). Consistent with these data, when these sparse NOE restraints were submitted to three independent structure calculation programs (CYANA, Xplor-NIH and ARIA) as part of a parallel methods study, the resulting calculations failed to converge on a common fold, with only 39 restraints forming a consensus across all three programs (Płoskoń et al., 2026).

**Figure 1:**
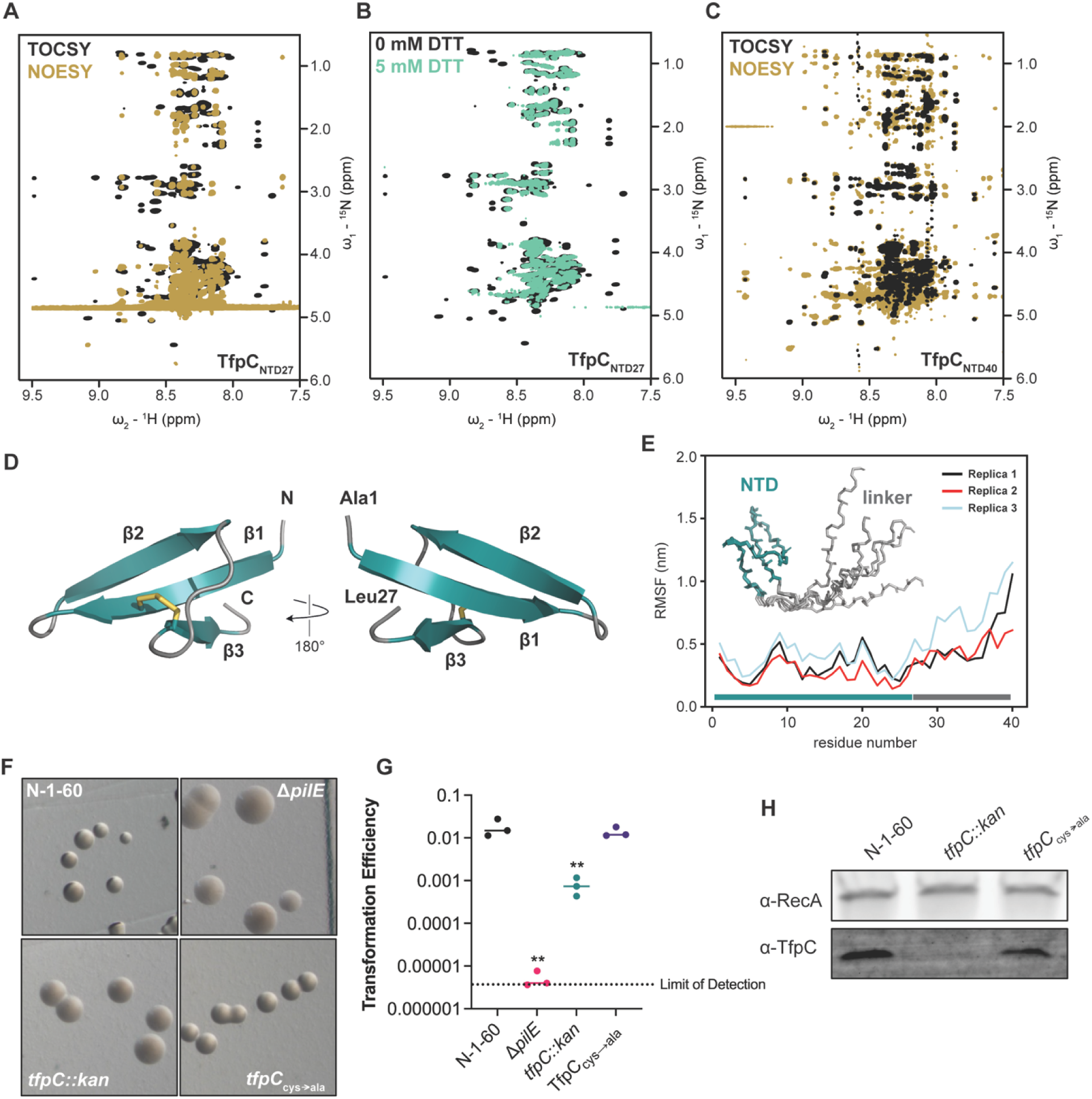
Structural characterization of the Gc TfpC N-terminal region. **A.** NMR ^1^H–^1^H TOCSY (black) and ^1^H–^1^H NOESY (gold) spectra of TfpC_NTD27_. **B.** NMR ^1^H–^1^H TOCSY spectra of TfpC_NTD27_ with (teal) and without (black) the addition of 5 mM DTT. **C.** NMR ^1^H–^1^H TOCSY (black) and ^1^H–^1^H NOESY (gold) spectra of TfpC_NTD40_. **D.** The top-ranked TfpC_NTD40_ Alphafold3 model showing residues 1-27. The disulfide bond between Cys6 and Cys22 is shown as yellow sticks. **E.** Mean RMSF of TfpC_NTD40_ C_α_ atoms taken from three independent simulations. Inset: top five ranked TfpC_NTD40_ Alphafold3 models. **F.** Gc colony morphology, imaged after 22 hours. **G.** Transformation efficiency assays. Efficiency is the number of nalidixic acid-resistant cells relative to the total number of cells plated. Statistical analysis was performed using a one-way ANOVA with Dunnett’s multiple comparisons to the parental strain. **, p = 0.0041; **, p = 0.0053. **H.** Western blot of Gc lysates. The anti-peptide TfpC antibody detects the C-terminus of TfpC. RecA is a loading control.

The N-terminus of TfpC contains two conserved cysteine residues, Cys6 and Cys22 (**Supplementary Figure 4**), and we speculated that these residues might form an intramolecular disulfide bond that would stabilize a domain in the oxidizing environment of the periplasm. In support of this, we observed additional, weaker resonances in spectra that showed greater dispersion in the amide proton region and were absent after adding dithiothreitol (5 mM, DTT) (**Figure 1B**). We reasoned that the initial peptide construct might be too short to support correct folding, and that an extended construct incorporating additional flanking sequence might increase the population of the folded state. We therefore analyzed an extended peptide comprising residues 1–40 (TfpC_NTD40_) (**Supplementary Figure 1**) and by NMR under the same experimental conditions, but this change did not improve the spectrum.

We next considered whether the population of the folded state might instead be temperature dependent. Although protein structures are often destabilized by heating, the hydrophobic effect that drives compaction of small polypeptides is known to strengthen with increasing temperature across the physiological range, favoring more compact, folded conformations before eventual thermal denaturation at much higher temperatures (Baldwin, 1986). Such compaction can, in turn, bring cysteine thiols into closer proximity and promote disulfide bond formation (Qin et al., 2015). We therefore increased the sample temperature from 4°C to 37°C, which markedly increased the relative intensity of the additional TOCSY resonances corresponding to the folded state and yielded substantially more long-range NOE correlations than were observed at the lower temperature (**Figure 1C**). Although spectral overlap between the two states prevented full assignment, we confidently assigned the chemical shifts of residues 6–15 in the folded state. The inter-residue NH-NH and NH-H_α_ NOE patterns were consistent with an antiparallel β-sheet, rather than the α-helix predicted by our previous model.

We used AlphaFold3 (Jumper et al., 2021) to model the structure of TfpC_NTD40_, which predicts a small, three-stranded antiparallel β-sheet (residues 2–25) followed by an extended region with variable structure (residues 26–40) (**Figure 1D, Supplementary Figure 5**). Within the folded domain, the β1 strand (residues 2–8) pairs with β2 (residues 11–16) via a tight β-turn at Asn9–Gly10, while a longer loop (residues 17–21) connects β2 to a third strand, β3 (residues 22–25). The β3 strand then packs against the opposite face of the β1 strand, and these strands are stapled via a disulfide bond between Cys6 and Cys22. Although the top five ranked AlphaFold3 models superimposed with little deviation across this β-core (residues 1-27; Root-Mean Squared Deviation (RMSD) of 0.21 ± 0.03 Å over core C_α_ atoms) and generated good Predicted Aligned Error (PAE) scores within this region (4.89 ± 0.41 Å), the predicted Local Distance Difference Test (pLDDT) scores were more modest (73.7 ± 1.4) (**Supplementary Figure 5**). We then ran a 200 ns molecular dynamics (MD) simulation (repeated at 180 ns) on the top-ranked TfpC_NTD40_ model, which remained stable throughout (**Figure 1E, Supplementary Figure 6**) and further supported the model’s validity. No clear structural homologs were identified for TfpC_NTD40_ using the DALI server (Holm et al., 2023) and this indicates that the TfpC N-terminal domain (NTD) forms a unique fold.

### The N-terminal disulfide bond is dispensable for pilus function

We next used genetic approaches to test the functional importance of the proposed NTD disulfide bond and the folded state that it stabilizes. To this end, we mutated both cysteine residues to alanine and measured the effect of this mutation on pilus function and TfpC protein stability. We qualitatively evaluate the piliation status of Gc strains using colony morphology assays (KELLOGG et al., 1963; Swanson et al., 1971) (**Figure 1F**). Colonies producing T4p, like our parental strain (N-1-60), are termed “piliated” and appear domed in shape with sharp borders. However, strains that do not produce T4p on the cell surface, like the Δ*pilE* strain, produce flat colonies with diffuse borders, termed “nonpiliated”. The *tfpC* mutant (*tfpC::kan*) produced colonies with an intermediate colony morphology phenotype, which we described previously as “underpiliated” (Hu et al., 2020). Mutating both TfpC cysteine residues to alanine (TfpC_cys→ala_) resulted in colony morphology similar to the parental strain.

Because the T4p mediates DNA uptake and natural transformation, we wanted to determine if TfpC’s two cysteine residues are important for transformation competence. The parental N-1-60 strain serves as a positive control for transformation efficiency, while the Δ*pilE* strain, which lacks the major pilus subunit, serves as a negative control and establishes the assay’s limit of detection (**Figure 1G**). As previously published, the *tfpC::kan* mutant shows significantly reduced transformation competence compared to the parental strain. Substituting the two TfpC cysteine residues with alanine residues retains transformation competence at levels equivalent to the parental strain. Consistent with the retention of transformation competence, the TfpC_cys→ala_ mutant did not impact TfpC protein abundance. Collectively, these genetic data show that TfpC’s potential N-terminal disulfide bond is dispensable for pilus function.

### Overall structure of the TfpC C-terminal domain

Our previous structural model of full-length TfpC (Hu et al., 2020) proposed a helical, coiled-coil (CC) domain at the C-terminus, which was supported by the presence of highly dispersed, downfield-shifted backbone amide resonances (¹H chemical shifts >8.5 ppm) in ¹H-¹ N HSQC spectra of full-length and N-terminally truncated (residues 52–147) TfpC. To define the boundaries of this domain more precisely and generate a construct suitable for structure determination, we analyzed the TfpC sequence using the Jpred4 server (Drozdetskiy et al., 2015), which predicted two helical regions spanning residues 86–110 and 120-144 of the 147-residue mature protein. A new recombinant C-terminally His_6_-tagged construct (TfpC_CC_, residues 81–147) was therefore created, and we elucidated its structure using solution heteronuclear NMR methods (**Supplementary Figure 1,7**).

From both manual and ARIA (Ambiguous Restraints for Iterative Assignment) NMR assignment methods, a total of 1819 NOEs were assigned in TfpC_CC_ ^15^N and ^13^C-edited NOESY spectra, and structure determination was also supplemented with Φ/φ dihedral angles and hydrogen bond restraints. The average pairwise RMSD for the water-refined final structures is 0.12 ± 0.03 Å for backbone atoms and 0.50 ± 0.12 Å for heavy atoms within secondary structure residues. Structural statistics are shown in **Supplementary Table 1**. TfpC_CC_ is composed of two antiparallel α-helices (residues 85–111 and 119–147) connected by a short turn. The two α-helices pack against each other along their length to form an elongated, rod-like coiled-coil hairpin approximately 50 Å in length (**Figure 2A,B**). The hydrophobic core is formed primarily by conserved residues Leu89, Leu93, Leu100, Ala103, Leu107 and Ala110 in helix 1, and Ile123, Leu126, Val130, Ile137, Leu140 and Leu144 in helix 2. The regular 3–4 residue spacing is characteristic of coiled-coil packing. Conserved Glu96 and Arg133 locally interrupt the otherwise continuous hydrophobic repeat, perturbing coiled-coil packing and introducing slight curvature into the otherwise extended two-helix assembly (**Supplementary Figure 8**). Although adjacent, Glu96 and Arg133 do not form a salt bridge. Instead, they lie within a cluster of acidic residues (Glu92, Glu102, Asp132), all of which are conserved except Glu102 (**Figure 2C**).

**Figure 2:**
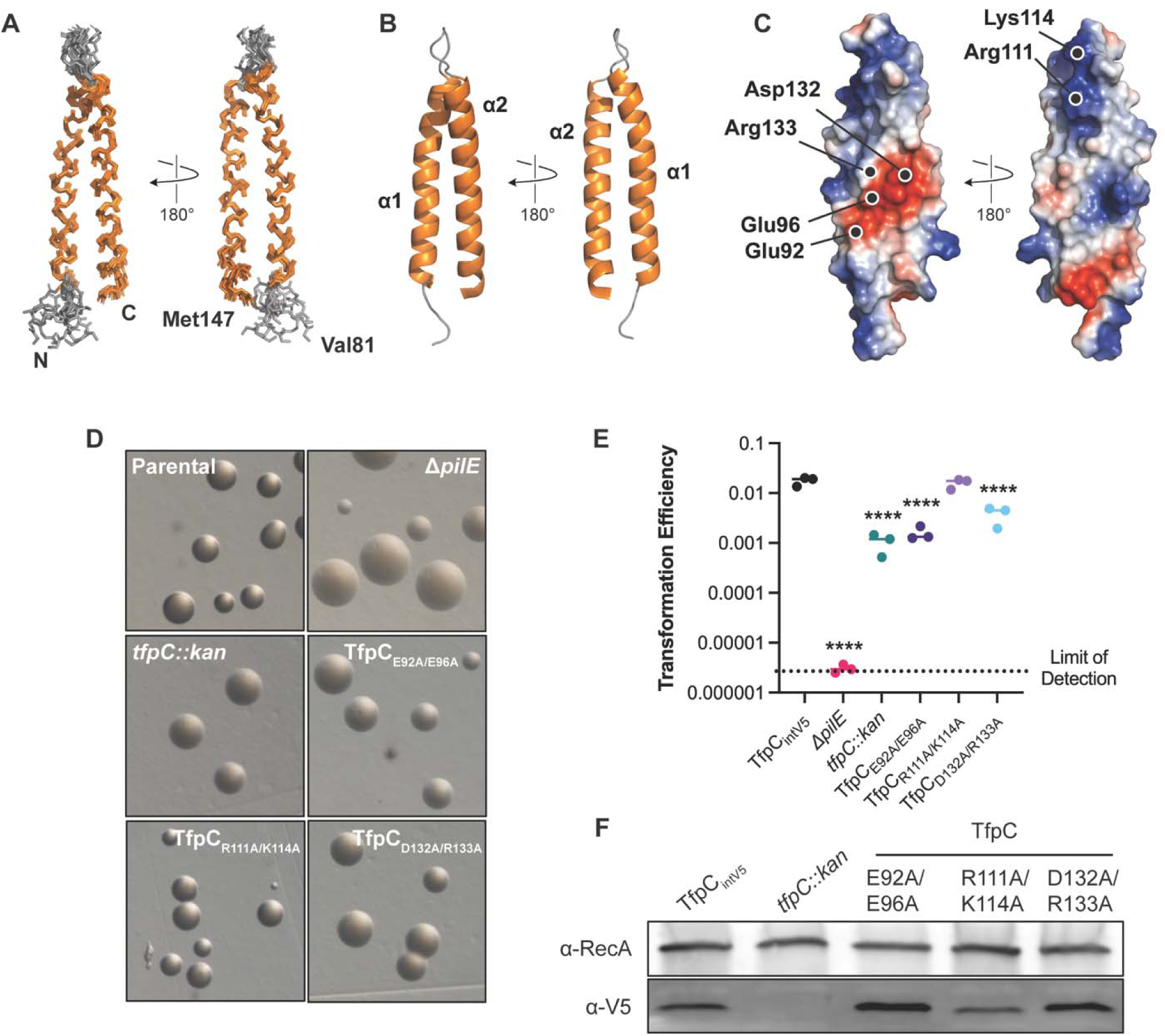
Structural characterization of the TfpC C-terminal region. **A.** Backbone superimposition of the ten best NMR structures of TfpC_CC_ shown in two orientations. **B.** Secondary structure of one of the final TfpC_CC_ structures shown as cartoon. **C.** Electrostatic surface potential of TfpC_CC_. **D.** Gc colony morphology, after 22 hours of growth. **E.** Transformation efficiencies, calculated by comparing the number of nalidixic-acid resistant cells relative to the total number of cells plated. Statistical analysis was performed using a one-way ANOVA with Dunnett’s multiple comparisons to the parental strain. ****, p < 0.0001. **F.** Western blot of Gc whole cell lysates (40 μg load) Anti-V5 detects epitope tagged TfpC. RecA is a loading control.

### The C-terminal domain of TfpC is important for pilus function

We next mutated three charged pairs to alanine to assess the importance of the conserved patch (Glu92/Glu96 and Asp132/Arg133) and, as a control, a basic patch (Arg111/Lys114) on the opposite face of the coiled-coil (**Figure 2C**). We named these mutants TfpC_E92A/E96A_, TfpC_D132A/R133A_, and TfpC_R111A/K114A_. We evaluated these strains for piliation qualitatively using colony morphology assays (**Figure 2D**). The *tfpC::kan* strain displayed an intermediate phenotype, illustrating loss of TfpC function. Similar to the TfpC loss-of-function mutant, both the TfpC_E92/E96A_ and TfpC_D132A/R133A_ mutants were underpiliated. However, the TfpC_R111A/K114A_ mutant showed pilus phenotypes similar to the parental strain.

We then assayed for T4p function using transformation efficiency assays, where the parental strain and Δ*pilE* strains served as positive and negative controls, respectively (**Figure 2E**). TfpC_E92A/E96A_ strains transformed similarly to the *tfpC* loss-of-function mutant, while the TfpC_R111A/K114A_ mutant transformed at parental levels. Interestingly, the TfpC_D132A/R133A_ mutant transformed at levels between the parental and *tfpC::kan* strains. We used western blotting to determine TfpC protein levels in each mutant, using the parental and *tfpC::kan* strains as controls (**Figure 2F**). Surprisingly, the TfpC_E92A/E96A_ and TfpC_D132A/R133A_ mutants produced TfpC protein at levels above the parental strain, while the TfpC_R111A/K114A_ mutant produced parental levels of TfpC. Together, these data suggest that mutations in the C-terminal acidic patch can impair TfpC function despite increasing TfpC protein levels. Replacement of Glu96 and Arg133 with alanine restores the hydrophobic character of the heptad repeat, potentially increasing coiled-coil stability. Thus, the native charged residues appear to contribute to TfpC function beyond maintaining the coiled-coil.

### The NTD or CTD alone are not sufficient for TfpC function

We wanted to evaluate the role of TfpC’s two structured domains by expressing these domains alone. However, because our TfpC antibody recognizes a C-terminal peptide epitope, we could not use it to detect truncated TfpC proteins lacking this region. We therefore generated a Gc strain expressing TfpC with an internal V5 tag, enabling detection of TfpC variants containing the tag (**Supplementary Figure 1**). We confirmed that internally tagging TfpC did not alter its function or stability (**Supplementary Figure 9**). For the N-terminal domain mutant where the N-terminus is expressed alone, we generated two versions—one with an N-terminal V5 tag between the signal sequence and the start of the mature protein (TfpC_NTD-NtV5_), and one with a C-terminal tag (TfpC_NTD-CtV5_; **Supplementary Figure 1**). For the C-terminal domain mutant expressed alone, we generated a version with an N-terminal V5 tag between the signal sequence and the start of the C-terminal domain (TfpC_CTD_; **Supplementary Figure 1**). Expressing either the N-or C-terminus alone phenocopied a TfpC mutant in colony morphology assays and transformation efficiency assays (**Figure 3A,B**). We ran western blots of whole-cell lysates and showed that neither the N-nor C-terminal domains were stably expressed (**Figure 3C**).

**Figure 3:**
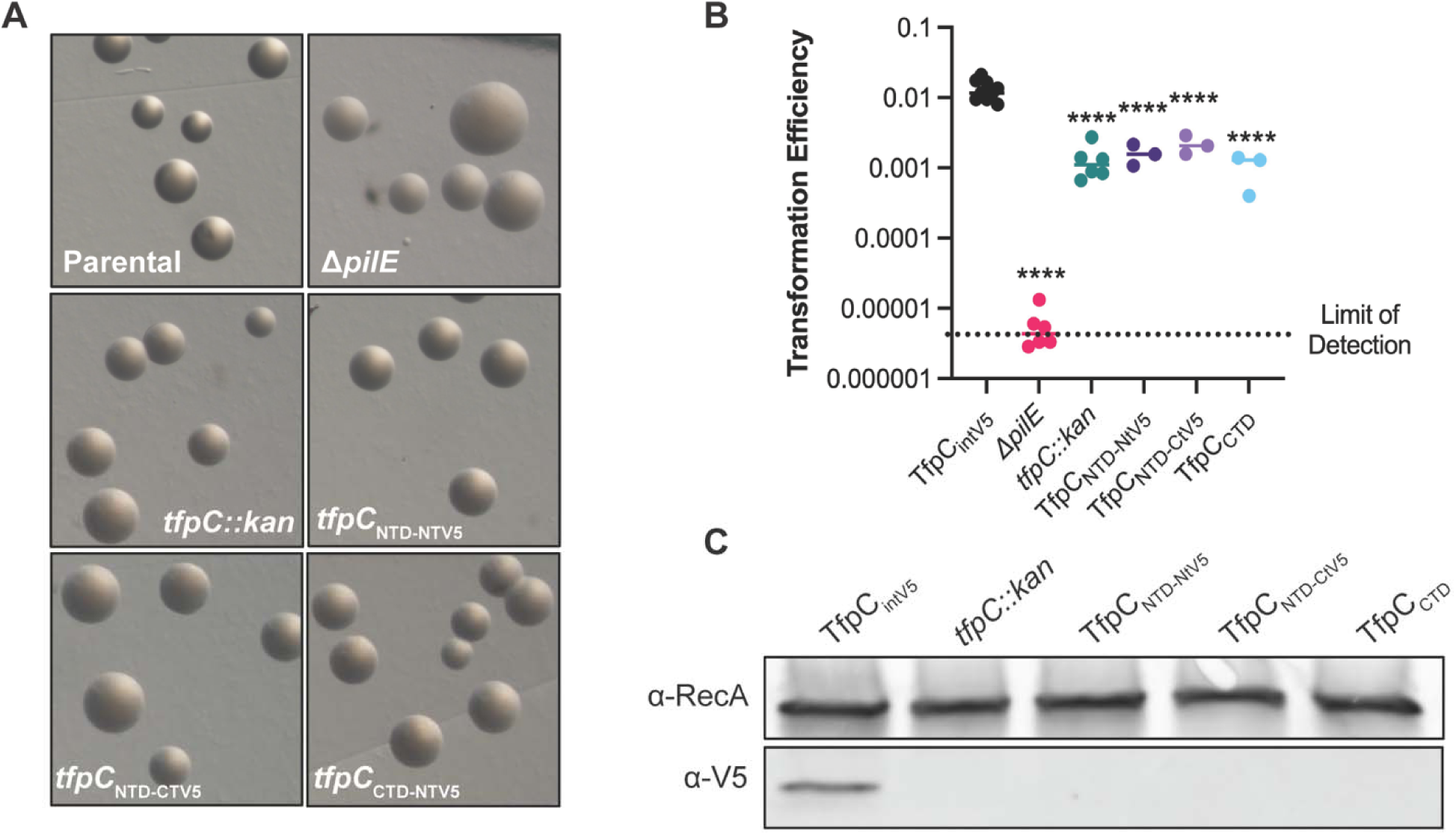
TfpC N-and C-terminal domains are both required for function. **A.** Colony morphology assays. **B.** Transformation efficiency of TfpC mutants, quantified by comparing the number of nalidixic acid-resistant cells to the total number of cells plated. Statistical analysis was performed using a one-way ANOVA with a Dunnett’s multiple comparisons test relative to the parental strain. ****, p < 0.0001. **C.** Western blot of Gc lysates detecting V5 epitope tagged TfpC. RecA is a loading control. 40 μg load.

### TfpC is a dynamic structure

TfpC contains a putative 53-amino acid disordered region between the N-and C-terminal domains (**Supplementary Figure 2**). Furthermore, this region contains 12 proline residues, and analysis using the PolyProline Predictor server (López-Sánchez et al., 2026) predicted that residues 41-51 (P^41^QTPEPAPSPS^51^) have the propensity to form a polyproline II (PPII) helix. To examine the global structure of full-length TfpC (TfpC_full_) in solution, we used small-angle X-ray scattering (SAXS) (**Supplementary Figure 1,10**, **Supplementary Table 2**). Guinier analysis suggested a radius of gyration (*R*_g_), the root mean square distance to the particles center of mass, of 3.56 nm, while analysis of the distance distribution function (P(r)) gave a maximum particle dimension (*D*_max_) of 19.5 nm and *R*_g_ of 4.36 nm. The difference between the Guinier-and (P(r))-derived *R*_g_ values is consistent with an elongated shape, as *R*_g_ reflects the distribution of mass around the particle’s centre of mass. Kratky plot analysis further supported this, indicating that TfpC is a highly dynamic particle in solution (Rambo & Tainer, 2011) (**Supplementary Figure 10**).

We therefore used the ensemble optimization method (EOM) to determine an ensemble of molecular models that best fit the SAXS data (Bernadó et al., 2007). As starting models, the NTD AlphaFold3 structure (residues 1–27) and the CTD NMR structure (residues 85–147) were used, connected by flexible linkers (**Figure 4A**). EOM analysis yielded good agreement between the experimental and calculated SAXS profiles (χ² = 2.8) (**Figure 4B**). The resulting ensemble comprised two approximately equal populations, with conformations spanning a continuum from more compact to more extended states (**Figure 4C,D, Supplementary Table 3**).

**Figure 4:**
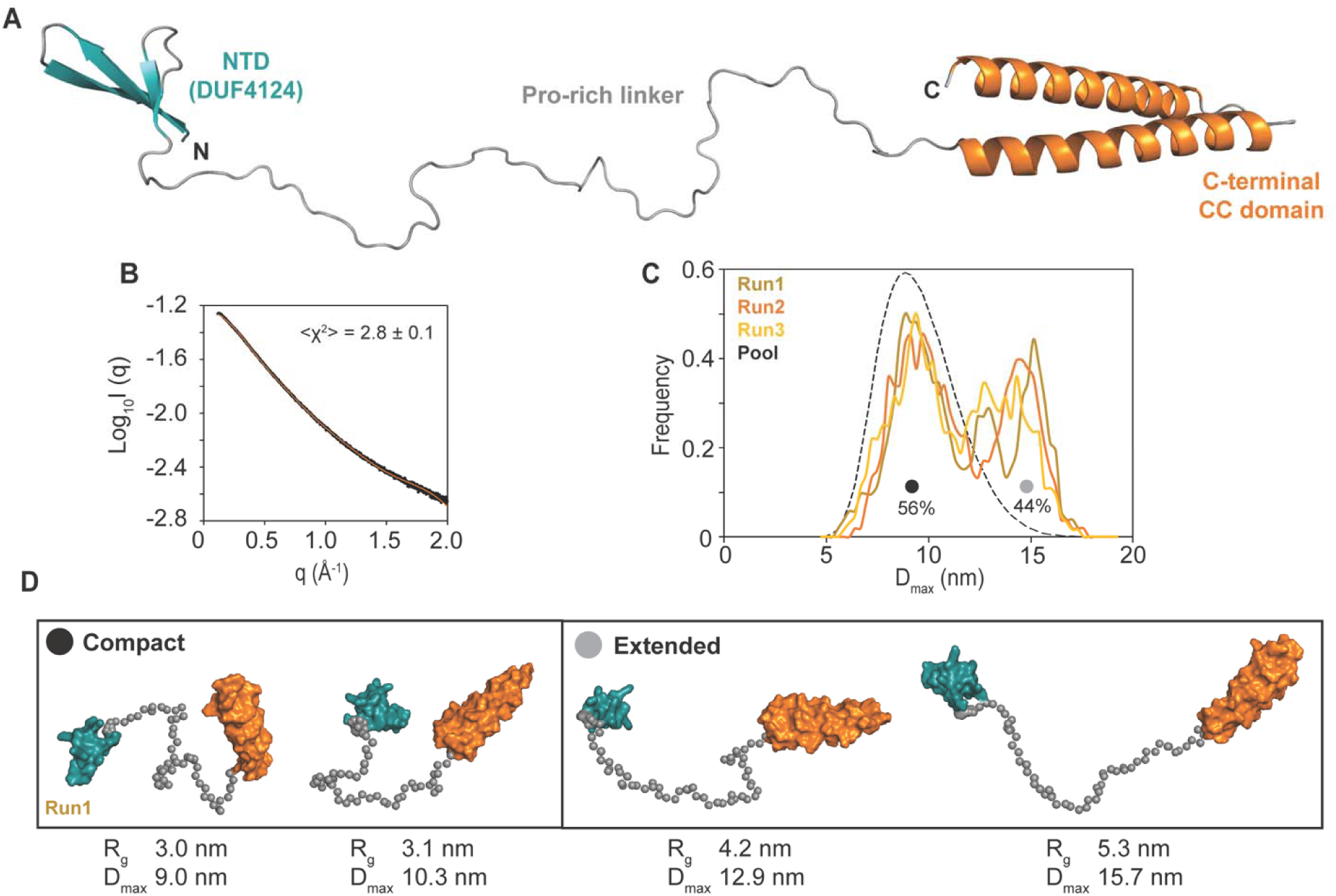
The TfpC disordered region is dispensable for pilus function, but not protein stability. **A.** Initial model of ChiA generated by EOM 2.0. **B.** EOM <FIT> (orange line) to the TfpC_full_ SAXS data (black open circles) with χ^2^ of 2.8. **C.** Three independent EOM runs yielded similar distributions of two populations. **D.** TfpC_full_ models corresponding to EOM run1 are shown.

### Mutations in the disordered region do not alter TfpC function

SAXS analysis showed that the TfpC linker is a dynamic region (DR) that allows substantial movement between the NTD and CTD. However, it did not reveal whether the linker adopts an ordered structure when TfpC interacts with other molecules or primarily acts as a spacer that maintains a defined distance between the two domains. To discern between these possibilities, we replaced 41-amino acids (residues 35-80 of the mature protein) from within the native disordered region with a new coding sequence of the same length (TfpC_newDR_; (**Supplementary Figure 1**) at the native *tfpC* genetic locus. We generated the random disordered sequence using the RandSeq server (Wilkins et al., 1999), excluding cysteine residues, and compared the disorder of intact TfpC and TfpC_newDR_ using AlphaFold3 (Jumper et al., 2021) to ensure that the modified linker retained a similar level of disorder. In addition, we generated mutants in which the disordered region was truncated by five (residues 52-56) and ten (residues 47-56) amino acids (TfpC_DRΔ5_ and TfpC_DRΔ10_; (**Supplementary Figure 1**). We measured the effect of these mutations on TfpC function using colony morphology assays and transformation efficiency assays (**Figure 5A,B**). The parental strain produced piliated colonies with high transformation efficiency, while the Δ*pilE* strain produced non-piliated, non-transformable colonies. The *tfpC::kan* mutant lacking TfpC protein produced underpiliated colonies with intermediate transformation efficiency. Like the parental strain, the TfpC_DRΔ5_, TfpC_DRΔ10_, and TfpC_newDR_ mutants also produced piliated colonies with high transformation efficiency. We next assessed TfpC protein levels by Western blot to determine whether the modified linker affected protein stability (**Figure 5C**). As expected, TfpC was detected in the parental strain but not in the *tfpC::kan* mutant. The TfpC_newDR_ strain produced lower levels of TfpC than the parental strain, while the TfpC_DRΔ5_ and TfpC_DRΔ10_ mutants similarly produced reduced TfpC levels. These data indicate that the disordered region of TfpC is dispensable for pilus function, but both its sequence and length are important for TfpC stability.

**Figure 5:**
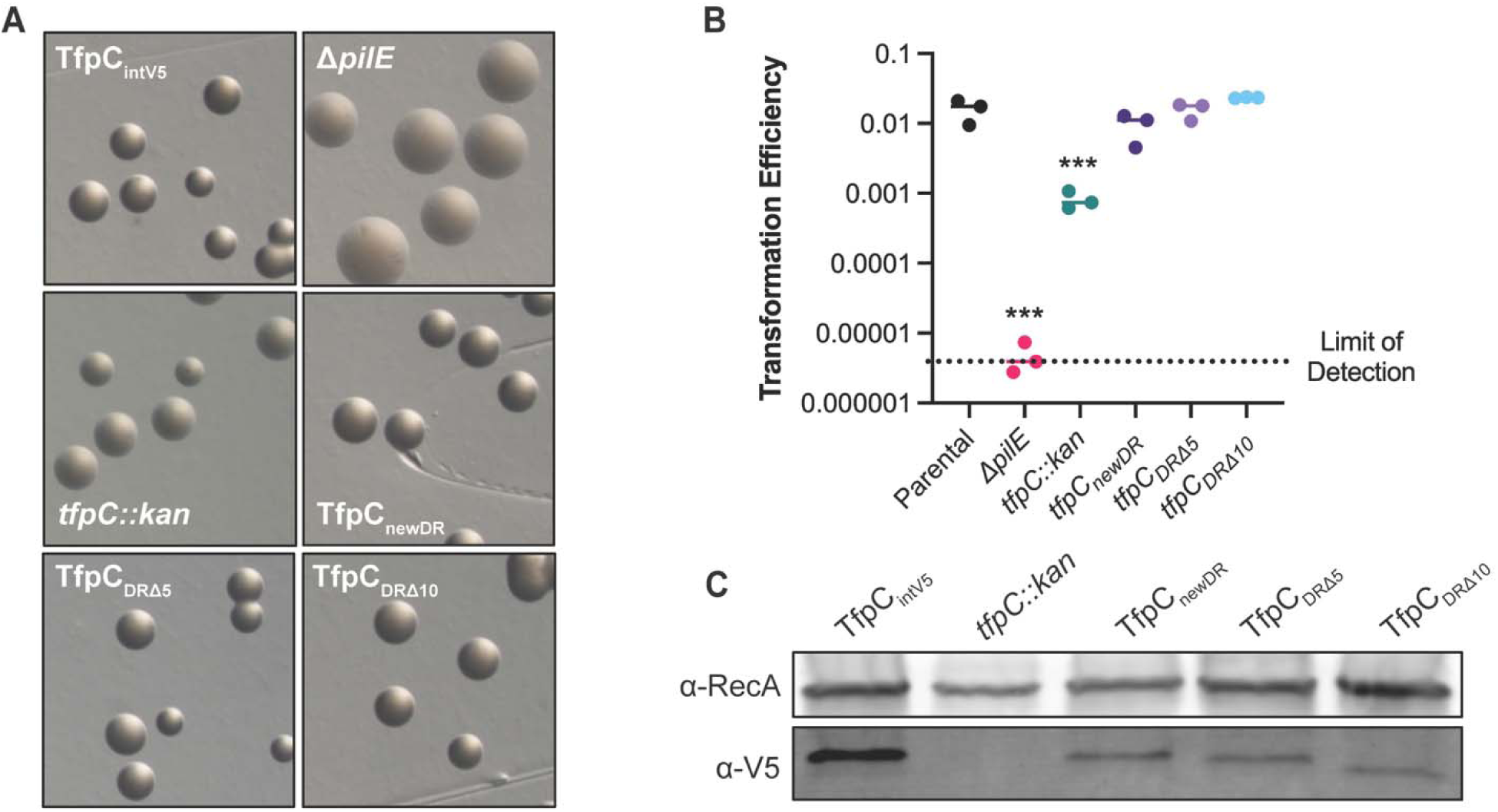
T**h**e **TfpC disordered region is dispensable for pilus function, but not protein stability. A.** Gc mutant colony morphology assays. **B.** Gc mutant transformation efficiency. Statistical analysis was performed using a one-way ANOVA with Dunnett’s multiple comparisons to the parental strain. ***, p = 0.0004; ***, p = 0.0007. **C.** Western blot of TfpC with 40 μg load. RecA serves as a loading control. V5 detects the TfpC protein.

## Discussion

TfpC is a periplasmic protein with three regions: a small N-terminal β-domain, a proline-rich disordered linker, and a C-terminal coiled-coil. Here, we characterized the domains separately and used genetic approaches to identify regions important for TfpC function and stability. Our findings allow us to assign structural and some functional features to each region.

The NTD is predominantly disordered but can adopt a novel three-stranded β-sheet, potentially stabilized by a disulfide bond between Cys6 and Cys22. The core of this structure is approximately 24 residues, which is at the lower size limit for an autonomously folding domain. Native folds of this size are usually disulfide-rich as they use these covalent bonds rather than a hydrophobic core to stabilize the fold (Cheek et al., 2006). However, mutating both cysteines to alanine had no detectable effect on piliation, transformation or TfpC abundance. One possibility is that the NTD forms a stable complex with another periplasmic component, reducing its need for disulfide stabilization.

The NTD fold belongs to the DUF4124 (Pfam PF13511) family (Hu et al., 2020), of which no experimentally determined structure has been reported to date. DUF4124-domain-containing proteins are found in a wide range of bacterial species and are often found with a variety of other domains. In most of these proteins, the DUF4124 domain forms the N-terminal region of the mature protein, upstream of a soluble lytic transglycosylase domain (Pfam PF01464). However, the same arrangement is also observed alongside M23 peptidase (Pfam PF01551), micrococcal nuclease (Pfam PF00565) and glutaredoxin (Pfam PF00462) domains. Although no binding partner has been reported for any DUF4124-domain protein, coupling of the DUF4124 domain with various enzymatic domains suggests that this fold is important for subcellular targeting and/or ligand binding. Since TfpC itself carries no predicted enzymatic domain, we predict it likely functions in periplasmic localization.

The C-terminal domain forms an antiparallel helical hairpin whose hydrophobic repeat is interrupted by Glu96 and Arg133, within an acidic face. Alanine mutation within this region (E92A/E96A, D132A/R133A) reduced piliation and increased TfpC protein levels, while mutation of a basic region on the opposing face (R111A/K114A) had no effect. The most likely explanation for the discrepancy between function and protein stability is that the acidic face of TfpC is a functionally important interaction surface, consistent with the roles of coiled-coil domains in other systems, which include oligomerization, membrane binding, molecular spacing, DNA recognition, and molecular scaffolding (Truebestein & Leonard, 2016). The increased TfpC stability found when replacing Glu96 and Arg133 with alanine could result from restoring the hydrophobic character of the coiled-coil heptad repeat, potentially increasing coiled-coil stability. An alternative explanation is that these changes increase protection of TfpC by an unknown binding partner.

The SAXS and bioinformatics analysis shows that the TfpC disordered region is highly flexible and may contain a short PPII helix. PPII helices are often observed within intrinsically disordered regions of proteins but are also known to mediate protein:protein interactions (Adzhubei et al., 2013). Disordered regions fall into eight categories, including entropic chains, which typically remain unstructured as part of their function, and can be further defined as flexible linkers and spacers (Trivedi & Nagarajaram, 2022). We found that the disordered region in the middle of TfpC can be replaced with a separate disordered region without completely destabilizing TfpC. Thus, it does not require sequence specificity to function, which is consistent with an entropic chain. Interestingly, shortening the disordered region by five or ten amino acids did not inactivate or fully destabilize TfpC, making this region less likely to be a spacer.

When the NTD or C-terminal domains were expressed in Gc, both proteins were unstable, which could indicate that both domains are required for stability. We predict that both the NTD and CTD interact with separate binding partners, and the disordered region is required to facilitate these independent binding activities. For disordered linkers this effect is determined by their length and dimensions rather than their sequence (Sørensen & Kjaergaard, 2019), which would also account for the reduced TfpC levels in our linker mutants, where the region remains disordered, but its length and composition are altered. Further studies are now needed to determine what cellular component(s) TfpC interacts with to facilitate pilus expression.

Here, we provide a detailed structural characterization of TfpC, a novel T4p component comprising three major domains. We show that the N-terminal DUF4124 domain can adopt a unique, dynamic β-sheet structure that is partially stabilized by disulfide-forming cysteine residues. The C-terminal domain forms an antiparallel helical hairpin containing an acidic patch required for TfpC function. These domains are connected via a proline-rich, disordered linker, which we hypothesize allows the N-or C-terminal domains to freely interact with their respective binding partners. This study combines structural and genetic approaches to define the structural and functional features of this previously unstudied protein family, providing a framework for understanding how TfpC contributes to type IV pilus function.

## Materials and Methods

### Bacterial Strains and Media

All gonococcal strains were derivatives of the FA1090 strain N-1-60 (Hu et al., 2020). In the N-1-60 strain, the guanine quadruplex site upstream of the *pilE* gene is mutated to prevent antigenic variation and the *pilC2* allele is locked in an “on” conformation to ensure expression of pili (Hu et al., 2020). Detailed strain information is listed in **Supplementary Table 4**. Gonococcal strains were grown on Gc medium base (GCB; Difco) (36.25 g/L), agar (1.25 g/L), Kellogg supplement I (22.2 mM glucose, 0.68 mM glutamine, 0.45 mM cocarboxylase) and II [1.23 mM Fe(NO_3_)_3_] (Dillard, 2011; Swanson et al., 1987).

### Generation of DNA Constructs

All gonococcal DNA constructs were purchased through Twist Biosciences (Twist Biosciences, South San Francisco, CA). All resulting mutants were verified using oligonucleotide primers purchased from Integrated DNA Technologies (IDT, Coralville, IA) and targeted Sanger DNA sequencing from GeneWiz. DNA corresponding to TfpC residues 81-147 (*tfpC_CC_*) was synthesized by Synbio Technologies and cloned into pET28b vector using NcoI and XhoI restriction sites. All constructs and oligonucleotides are listed in **Supplementary Table 5**, and synthesised genes are listed in **Supplementary Table 6**.

### Protein purification

Intact TfpC (residues 1 to 147) was expressed as described previously (Hu et al., 2020). Expression was carried out in *E. coli* BL21(DE3) (New England Biolabs) grown in LB media in the presence of 50 μg/ml kanamycin at 37°C. Expression was induced with 0.5□mM isopropyl-d-1-thiogalactopyranoside (IPTG) at an OD_600_ of 0.6, and cells were harvested after growth overnight at 18°C. Cells were resuspended in 20□mM Tris-HCl (pH 8), 200□mM NaCl, lysed by sonication, and purified using nickel affinity chromatography (New England Biolabs). Samples were then gel filtered using a Superdex 200 column (Cytiva) equilibrated in 20 mM Tris-HCl (pH 8), 200 mM NaCl. The TfpC coiled-coil domain (TfpC_81-147_, residues 81-147) was expressed in *E. coli* BL21(DE3) (New England Biolabs), where cells were grown in the presence of 50 μg/ml kanamycin at 37°C in M9 minimal medium containing 0.07% (w/v) ^15^NH_4_Cl_2_ and 0.2% (w/v) [^13^C]glucose (Cambridge Isotope Laboratories). Expression was induced with 0.5 mM IPTG at an OD_600_ of 0.6, and cells were harvested after growth overnight at 18°C. Cells were resuspended in 20 mM Tris-HCl (pH 8), 200 mM NaCl, lysed by sonication, and purified using nickel affinity chromatography (New England Biolabs). Samples were then gel filtered using a Superdex 75 column (Cytiva) equilibrated in 50 mM sodium phosphate buffer (pH 7.5), 100 mM NaCl.

### Peptide NMR

All peptides were synthesised by Genscript to >95% purity. TfpC_1-27_ peptide (AKIYTCTINGETVYTTKPSKSCHSTDL-NH_2_) was resuspended at 3 mM, and TfpC_1-40_ peptide (AKIYTCTINGETVYTTKPSKSCHSTDLPPIGNYSSERYIL-NH_2_) was resuspended at 5 mM, in 50 mM sodium acetate (D_₃_, 99%; pH 5.5), 50 mM NaCl, 10% or 100% (v/v) D_2_O. Backbone and side chain assignments were achieved using standard double-resonance peptide assignment experiments (^1^H-^15^N heteronuclear single quantum coherence (HSQC), ^1^H-^1^H Total Correlation Spectroscopy (TOCSY), ^1^H-^1^H correlation spectroscopy (COSY), ^1^H-^1^H Nuclear Overhauser Effect Spectroscopy (NOESY) (τ_m_: 300 ms, TfpC_1-27_; 120 ms, TfpC_1-40_), ^1^H-^13^C HSQC, ^1^H-^13^C Heteronuclear Multiple Bond Correlation (HMBC), ^1^H-^13^C heteronuclear multiple quantum coherence (HMQC)-TOCSY) recorded at temperatures between 278 K and 310 K on 700, 800 MHz Bruker Avance IIIHD and 950 MHz Bruker Avance Neo NMR spectrometers, equipped with cryoprobes. All spectra were processed using NMRPipe (Delaglio et al., 1995) and analysed using CCPNMRv3 (Skinner et al., 2016).

### Heteronuclear NMR

Measurements were performed at 298 K on a ^15^N^13^C-labelled TfpC_81-147_ sample (0.25 mM) in 50 mM sodium phosphate buffer (pH 7.5), 100 mM NaCl, 10% D_2_O on 700, 800 MHz Bruker Avance IIIHD and 950 MHz Bruker Neo NMR spectrometers, equipped with cryoprobes. Assignments were completed using standard triple-resonance assignment methodology (Sattler, 1999). A total of 86% of the native TfpC_81-147_ backbone (e.g., disregarding proline residues) and 94% of the native TfpC_81-147_ side-chain resonances were assigned. NOE data for structure calculations of TfpC_81-147_ were obtained from 800-MHz edited ^1^H-^15^N and 950-MHz edited ^1^H-^13^C 3D NOESY-heteronuclear single/multiple quantum coherence (HSQC/HMQC) experiments. The ARIA protocol for automated NOESY assignment interfaced with the CNS program was used for structure calculation through the ARIAweb server (Allain et al., 2020). Secondary structure was first identified using the chemical shift-based dihedral angle prediction software TALOS (Cornilescu et al., 1999). For residues located in secondary structure, hydrogen bonds and /φ backbone dihedral angles derived from TALOS were introduced as restraints in the ARIA structure calculation. The frequency window tolerances for assigning NOEs were ±0.02 ppm for direct proton dimensions, ±0.04 ppm for indirect proton dimensions, and ±0.50 ppm for nitrogen and carbon dimensions. Fifty structures were calculated in the final iteration, and these had no NOE violations >0.3 Å and no dihedral angle violations >5°. All spectra were processed using NMRPipe (Delaglio et al., 1995) and analysed using CCPNMRv3 (Skinner et al., 2016). A summary of NMR-derived restraints and statistics is reported in **Supplementary Table 1**.

### Molecular Dynamics

The top ranking AlphaFold3 model of TfpC_NTD40_ was run through the CHARMM-GUI (Jo et al., 2008) solution builder server and placed in 0.15 M NaCl solution with 75.2 × 75.2 × 75.2 Å^3^ box dimension, resulting in a total of 80 salt, 14078 water, and 619 protein atoms. MD simulations were run using GROMACS 2019 (Abraham et al., 2015) and the CHARMM36 force field (Huang et al., 2017). The system was thermalized, equilibrated, and simulated at 303.5 K and 1 bar pressure following the simulation protocol suggested by CHARMM-GUI (Jo et al., 2008). From the final structure after pressure equilibration, one independent production trajectories of 200 ns and two independent production trajectories of 180 ns, that resulted in statistically consistent configurational ensembles, were generated. Trajectories were analysed using the python package MDAnalysis (Michaud-Agrawal et al., 2011).

### SEC-SAXS

Data were collected on beamline at the Diamond Light Source (DLS), UK (Cowieson et al., 2020). 60 μl of TfpC_full_ (25 mg/ml) in 20 mM Tris–HCl pH 8, 200 mM NaCl was applied to a Shodex KW403-4F column at 0.16 ml/min and SAXS data were measured over a momentum transfer range of 0.004<*q*<0.34 Å^−1^. Peak integration and buffer subtraction were performed in CHROMIXS (Panjkovich & Svergun, 2018). The radius of gyration (*R*_g_) and scattering at zero angle (I(0)) were calculated from the analysis of the Guinier region by AUTORG (Franke et al., 2017). The distance distribution function (*P*(r)) was subsequently obtained using GNOM (Franke et al., 2017), yielding the maximum particle dimension (*D*_max_). Determination of molecular model ensembles that best fit the SAXS data was performed in triplicate using EOM2 (Tria et al., 2015). SAXS structural and EOM parameters can be found in **Supplementary Table 2,3**.

### Imaging of Pilus-Dependent Colony Morphology

*N. gonorrhoeae* colonies grown for 22 hours on a thin layer of solid GCB medium (20 ml per 100 x 100 x 15 mm plate, Alkali Scientific, Fort Lauderdale, FL) were observed and photos were taken using a Nikon SMZ-10A stereomicroscope and a Nikon digital-site camera.

### Transformation Efficiency Assays

Transformation assays were performed similarly as previously published (Hu et al., 2020; Nicholson et al., 2025). Gc was struck from frozen stocks and grown overnight on GCB plates at 37°C under 5% CO_2_. Gc was collected in 400 μl of GCBL (supplemented with Kellogg supplement I and II and 5 mM MgSO_4_) using a polyester swab. Cultures were diluted to an OD_550_ of 0.15 in a 200 μl total volume of GCBL. 2 μl of a 25 ng/μl pSY6 plasmid (Stein et al., 1991) DNA stock was added to each 200 μl reaction and incubated at 37°C for 20 min. Then, 1 U of DNase I was added to each tube and incubated at 37°C for 10 min. For each reaction, the entire reaction was then added to 2 ml of 37°C prewarmed GCBL media with no compound or 1 μM SC144 in a 24-well plate and incubated at 37°C under 5% CO_2_ for 2 h. These 2.2 ml reactions were mixed, and a portion was harvested for 10-fold serial dilutions for CFU plating. Serial dilutions were performed using GCBL. 10 μl of each serial dilution from 10^0^ to 10^-6^ was plated on both GCB and GCB containing 1 μg/ml nalidixic acid. CFU were counted using a stereoscope after overnight growth at 37°C under 5% CO_2_.

### Western Blotting

Gc was struck from frozen stocks and grown overnight on GCB plates at 37°C under 5% CO_2_. Single colonies were picked using a pipette tip and heavy struck on GCB plates. These heavy streaks were grown overnight at 37°C under 5% CO_2_. The entire plate of Gc was harvested in 450 μl of 1X PBS containing 0.1% phenylmethylsulfonyl fluoride (PMSF, Roche) using a polyester swab. Total protein was quantified using the Pierce BCA Protein Assay kit (ThermoFisher). 40 μg of protein was mixed with 5X sample buffer containing β-mercaptoethanol and boiled for 10 minutes. Samples were run on 4-15% Tris-glycine SDS-PAGE gels (Bio-Rad). Gels were transferred to methanol-activated PVDF membrane and blocked overnight at 4°C in 5% milk in 1X TBS + 0.1% Tween-20. The α-TfpC antibody (Pacific Immunology, Ramona, CA, USA) detects the DRQQNIQALQRELGRM epitope at the C-terminus of TfpC. Blots were incubated with the TfpC antibody (1:500 dilution in 5% milk in TBS-T) overnight at 4°C. For the α-V5 antibody (ThermoFisher) blots were incubated in a 1:5,000 dilution (5% milk in TBS-T) overnight at 4°C. To detect RecA, a polyclonal *E. coli* α-RecA antisera (a generous gift from Mike Cox) was used at 1:2,000 (in 5% milk in TBS-T) for one hour at room temperature. Secondary antibodies, Goat anti-Rabbit CyDye 800 (Cytiva) and Goat anti-Mouse IgG (H+L) Alexa Fluor 700 (Invitrogen), were used at 1:20,000 dilutions in 5% milk in TBS-T for two hours at room temperature. The Goat anti-Chicken IgY (H+L) Alexa Fluor 488 (Invitrogen) antibody was used at a 1:10,000 dilution. Western blots were imaged on an Amersham Typhoon.

### Multiple Sequence Alignments and Visualization

We used NCBI BLASTP to identify and align sequences to the TfpC amino acid sequence (Altschul et al., 1997). We performed this BLASTP (version 2.17.0+) under default parameters (ClusteredNR database), searching within *Neisseria* (taxid:482). Using the BLASTp output and the NCBI Constraint-based Multiple Alignment Tool (COBALT), we generated an alignment file (with gaps) containing the top 100 hits. Sequence alignments were visualized in the ESPript 3 web server (Gouet et al., 1999).

## Statistical Analysis

All statistical analyses were performed in GraphPad Prism version 10. Statistical significance was determined by performing a One-Way ANOVA followed by a Dunnett’s multiple comparison’s test. P-values for individual experiments are noted in the figure legends. All experiments were conducted on at least three biological replicates, each with technical triplicates where available.

## Data Availability

The assigned chemical shifts of TfpC_CC_ have been deposited in the BioMagResBank (http://www.bmrb.wisc.edu/) with accession code XXXXX. Coordinates for TfpC_CC_ have been deposited in the PDB (https://www.rcsb.org/) with accession code XXXX. The buffer subtracted SAXS curves for TfpC_full_ have been deposited in the Small Angle Scattering Biological Data Bank (https://www.sasbdb.org) with accession ID SASDXXX.

## Acknowledgements.

This work was supported by MRC grants (MR/W000814/1 and UKRI1453) awarded to J.G. M.B. was supported by a BBSRC LIDO PhD studentship (BB/T008709/1) awarded to J.G. and C.L. This work was also supported by the Francis Crick Institute through provision of access to the MRC Biomedical NMR Centre. The Francis Crick Institute receives its core funding from Cancer Research UK (FC001029), the UK Medical Research Council (CC1078), and the Wellcome Trust (CC1078). Research reported in this publication was supported by the National Institute of Allergy and Infectious Disease (NIAID) of the National Institutes of Health under grant number R01AI146073 to H.S.S. K.R.N was supported by an Immunology and Molecular Pathogenesis Training Program fellowship T32 (T32AI007476) and a Ruth L. Kirschstein NRSA F32 (F32AI186488). The content is solely the responsibility of the authors and does not necessarily represent the official views of the National Institutes of Health. This manuscript is the result of funding in whole or in part by the National Institutes of Health (NIH). It is subject to the NIH Public Access Policy. Through acceptance of this federal funding, NIH has been given a right to make this manuscript publicly available in PubMed Central upon the Official Date of Publication, as defined by NIH. This work made use of time on HPC granted via the UK High-End Computing Consortium for Biomolecular Simulation, HECBioSim (http://hecbiosim.ac.uk), supported by EPSRC (grant no. EP/R029407/1). We also acknowledge the King’s Computational Research, Engineering and Technology Environment (CREATE) at King’s College London for providing us access to computational resources. We thank the beamline scientists at BL21 of the Diamond Light Source, United Kingdom.

## Author contributions

Conceived and designed the experiments: KN, MB, EP-A, PS, AO, CL, HS, JG. Performed the experiments: KN, MB, EP-A, PS, AO, JG. Analyzed the data: KN, MB, EP-A, PS, AO, CL, HS, JG. Contributed reagents/materials/analysis tools: CL, HS, JG. Wrote the paper: KN, MB, EP-A, PS, AO, CL, HS, JG.

## Additional information

Competing interests. The authors declare no competing interests.

## Notes

### Competing Interest Statement

The authors have declared no competing interest.

## References

Abraham, M. J., Murtola, T., Schulz, R., Páll, S., Smith, J. C., Hess, B., & Lindahl, E. (2015). GROMACS: High performance molecular simulations through multi-level parallelism from laptops to supercomputers. SoftwareX, *1–2*, 19–25. 10.1016/j.softx.2015.06.001

Adzhubei, A. A., Sternberg, M. J. E., & Makarov, A. A. (2013). Polyproline-II helix in proteins: structure and function. Journal of Molecular Biology, 425(12), 2100–2132. 10.1016/j.jmb.2013.03.018

Allain, F., Mareuil, F., Ménager, H., Nilges, M., & Bardiaux, B. (2020). ARIAweb: a server for automated NMR structure calculation. Nucleic Acids Research, 48(W1), W41–W47. 10.1093/nar/gkaa362

Altschul, S. F., Madden, T. L., Schäffer, A. A., Zhang, J., Zhang, Z., Miller, W., & Lipman, D. J. (1997). Gapped BLAST and PSI-BLAST: a new generation of protein database search programs. Nucleic Acids Research, 25(17), 3389–3402. 10.1093/nar/25.17.3389

Baldwin, R. L. (1986). Temperature dependence of the hydrophobic interaction in protein folding. Proceedings of the National Academy of Sciences of the United States of America, 83(21), 8069–8072. 10.1073/pnas.83.21.8069

Bernadó, P., Mylonas, E., Petoukhov, M. V, Blackledge, M., & Svergun, D. I. (2007). Structural characterization of flexible proteins using small-angle X-ray scattering. Journal of the American Chemical Society, 129(17), 5656–5664. 10.1021/ja069124n

Berry, J.-L., Phelan, M. M., Collins, R. F., Adomavicius, T., Tønjum, T., Frye, S. A., Bird, L., Owens, R., Ford, R. C., Lian, L.-Y., & Derrick, J. P. (2012). Structure and assembly of a trans-periplasmic channel for type IV pili in Neisseria meningitidis. PLoS Pathogens, 8(9), e1002923. 10.1371/journal.ppat.1002923

Carbonnelle, E., Hélaine, S., Prouvensier, L., Nassif, X., & Pelicic, V. (2005). Type IV pilus biogenesis in Neisseria meningitidis: PilW is involved in a step occurring after pilus assembly, essential for fibre stability and function. Molecular Microbiology, 55(1), 54–64. 10.1111/j.1365-2958.2004.04364.x

Cehovin, A., Simpson, P. J., McDowell, M. A., Brown, D. R., Noschese, R., Pallett, M., Brady, J., Baldwin, G. S., Lea, S. M., Matthews, S. J., & Pelicic, V. (2013). Specific DNA recognition mediated by a type IV pilin. Proceedings of the National Academy of Sciences of the United States of America, 110(8), 3065–3070. 10.1073/pnas.1218832110

Chang, Y.-W., Rettberg, L. A., Treuner-Lange, A., Iwasa, J., Søgaard-Andersen, L., & Jensen, G. J. (2016). Architecture of the type IVa pilus machine. *Science (New York*, N.Y.), 351(6278), aad2001. 10.1126/science.aad2001

Cheek, S., Krishna, S. S., & Grishin, N. V. (2006). Structural classification of small, disulfide-rich protein domains. Journal of Molecular Biology, 359(1), 215–237. 10.1016/j.jmb.2006.03.017

Cornilescu, G., Delaglio, F., & Bax, A. (1999). Protein backbone angle restraints from searching a database for chemical shift and sequence homology. Journal of Biomolecular NMR, 13(3), 289–302. 10.1023/a:1008392405740

Cowieson, N. P., Edwards-Gayle, C. J. C., Inoue, K., Khunti, N. S., Doutch, J., Williams, E., Daniels, S., Preece, G., Krumpa, N. A., Sutter, J. P., Tully, M. D., Terrill, N. J., & Rambo, R. P. (2020). Beamline B21: high-throughput small-angle X-ray scattering at Diamond Light Source. Journal of Synchrotron Radiation, 27(Pt 5), 1438–1446. 10.1107/S1600577520009960

Craig, L., Forest, K. T., & Maier, B. (2019). Type IV pili: dynamics, biophysics and functional consequences. Nature Reviews. Microbiology, 17(7), 429–440. 10.1038/s41579-019-0195-4

Craig, L., & Li, J. (2008). Type IV pili: paradoxes in form and function. Current Opinion in Structural Biology, 18(2), 267–277. 10.1016/j.sbi.2007.12.009

Delaglio, F., Grzesiek, S., Vuister, G. W., Zhu, G., Pfeifer, J., & Bax, A. (1995). NMRPipe: a multidimensional spectral processing system based on UNIX pipes. Journal of Biomolecular NMR, 6(3), 277–293. 10.1007/BF00197809

Dillard, J. P. (2011). Genetic Manipulation of Neisseria gonorrhoeae. *Current Protocols in Microbiology*, Chapter 4, Unit4A.2. 10.1002/9780471729259.mc04a02s23

Drozdetskiy, A., Cole, C., Procter, J., & Barton, G. J. (2015). JPred4: a protein secondary structure prediction server. Nucleic Acids Research, 43(W1), W389–94. 10.1093/nar/gkv332

Franke, D., Petoukhov, M. V, Konarev, P. V, Panjkovich, A., Tuukkanen, A., Mertens, H. D. T., Kikhney, A. G., Hajizadeh, N. R., Franklin, J. M., Jeffries, C. M., & Svergun, D. I. (2017). ATSAS 2.8: a comprehensive data analysis suite for small-angle scattering from macromolecular solutions. Journal of Applied Crystallography, 50(Pt 4), 1212–1225. 10.1107/S1600576717007786

Georgiadou, M., Castagnini, M., Karimova, G., Ladant, D., & Pelicic, V. (2012). Large-scale study of the interactions between proteins involved in type IV pilus biology in Neisseria meningitidis: characterization of a subcomplex involved in pilus assembly. Molecular Microbiology, 84(5), 857–873. 10.1111/j.1365-2958.2012.08062.x

Gouet, P., Courcelle, E., Stuart, D. I., & Metoz, F. (1999). ESPript: analysis of multiple sequence alignments in PostScript. Bioinformatics, 15(4), 305–308. 10.1093/bioinformatics/15.4.305

Helaine, S., Dyer, D. H., Nassif, X., Pelicic, V., & Forest, K. T. (2007). 3D structure/function analysis of PilX reveals how minor pilins can modulate the virulence properties of type IV pili. Proceedings of the National Academy of Sciences of the United States of America, 104(40), 15888–15893. 10.1073/pnas.0707581104

Holm, L., Laiho, A., Törönen, P., & Salgado, M. (2023). DALI shines a light on remote homologs: One hundred discoveries. Protein Science: A Publication of the Protein Society, 32(1), e4519. 10.1002/pro.4519

Hu, L. I., Yin, S., Ozer, E. A., Sewell, L., Rehman, S., Garnett, J. A., & Seifert, H. S. (2020). Discovery of a New Neisseria gonorrhoeae Type IV Pilus Assembly Factor, TfpC. mBio, 11(5). 10.1128/mBio.02528-20

Huang, J., Rauscher, S., Nawrocki, G., Ran, T., Feig, M., de Groot, B. L., Grubmüller, H., & MacKerell, A. D. (2017). CHARMM36m: an improved force field for folded and intrinsically disordered proteins. Nature Methods, 14(1), 71–73. 10.1038/nmeth.4067

Jo, S., Kim, T., Iyer, V. G., & Im, W. (2008). CHARMM-GUI: a web-based graphical user interface for CHARMM. Journal of Computational Chemistry, 29(11), 1859–1865. 10.1002/jcc.20945

Jumper, J., Evans, R., Pritzel, A., Green, T., Figurnov, M., Ronneberger, O., Tunyasuvunakool, K., Bates, R., Žídek, A., Potapenko, A., Bridgland, A., Meyer, C., Kohl, S. A. A., Ballard, A. J., Cowie, A., Romera-Paredes, B., Nikolov, S., Jain, R., Adler, J.,…Hassabis, D. (2021). Highly accurate protein structure prediction with AlphaFold. Nature, 596(7873), 583–589. 10.1038/s41586-021-03819-2

Kellogg, D. S., Peacock, W. L., Deacon, W. E., Brown, L., & Pirkle, D. I. (1963). NEISSERIA GONORRHOEAE. I. VIRULENCE GENETICALLY LINKED TO CLONAL VARIATION. Journal of Bacteriology, 85(6), 1274–1279. 10.1128/jb.85.6.1274-1279.1963

Kolappan, S., Coureuil, M., Yu, X., Nassif, X., Egelman, E. H., & Craig, L. (2016). Structure of the Neisseria meningitidis Type IV pilus. Nature Communications, 7, 13015. 10.1038/ncomms13015

López-Sánchez, R., Pantoja-Uceda, D., Mompeán, M., & Laurents, D. V. (2026). PolyProline Predictor: A web server for empirical sequence-based prediction of polyproline II helices. Protein Science: A Publication of the Protein Society, 35(7), e70675. 10.1002/pro.70675

Merz, A. J., & So, M. (2000). Interactions of pathogenic neisseriae with epithelial cell membranes. Annual Review of Cell and Developmental Biology, 16, 423–457. 10.1146/annurev.cellbio.16.1.423

Merz, A. J., So, M., & Sheetz, M. P. (2000). Pilus retraction powers bacterial twitching motility. Nature, 407(6800), 98–102. 10.1038/35024105

Michaud-Agrawal, N., Denning, E. J., Woolf, T. B., & Beckstein, O. (2011). MDAnalysis: a toolkit for the analysis of molecular dynamics simulations. Journal of Computational Chemistry, 32(10), 2319–2327. 10.1002/jcc.21787

Muir, A., Gurung, I., Cehovin, A., Bazin, A., Vallenet, D., & Pelicic, V. (2020). Construction of a complete set of Neisseria meningitidis mutants and its use for the phenotypic profiling of this human pathogen. Nature Communications, 11(1), 5541. 10.1038/s41467-020-19347-y

Nicholson, K. R., Yin, S., Edwards, J. L., Luan, C.-H., & Seifert, H. S. (2025). Natural compounds target the M23B zinc metallopeptidase Mpg to modulate Neisseria gonorrhoeae Type IV pilus expression. mBio, 16(4), e0402724. 10.1128/mbio.04027-24

Obergfell, K. P., & Seifert, H. S. (2016). The Pilin N-terminal Domain Maintains Neisseria gonorrhoeae Transformation Competence during Pilus Phase Variation. PLoS Genetics, 12(5), e1006069. 10.1371/journal.pgen.1006069

Panjkovich, A., & Svergun, D. I. (2018). CHROMIXS: automatic and interactive analysis of chromatography-coupled small-angle X-ray scattering data. *Bioinformatics (Oxford*, England), 34(11), 1944–1946. 10.1093/bioinformatics/btx846

Płoskoń, E., Baskaran, K., Tejero, R., Schwieters, C. D., Bardiaux, B., Guentert, P., Fogh, R. H., Gutmanas, A., Brooksbank, E. J., Yokochi, M., Wishart, D. S., Wedell, J. R., Vranken, W. F., Thompson, D., Thomson, G., Smith, B. O., Rehman, S., Ramelot, T. A., Ragan, T. J.,…Vuister, G. W. (2026). The NMR Exchange Format (NEF): Specification and Applications. bioRxiv: The Preprint Server for Biology. 10.64898/2026.04.22.715536

Qin, M., Wang, W., & Thirumalai, D. (2015). Protein folding guides disulfide bond formation. Proceedings of the National Academy of Sciences of the United States of America, 112(36), 11241–11246. 10.1073/pnas.1503909112

Rambo, R. P., & Tainer, J. A. (2011). Characterizing flexible and intrinsically unstructured biological macromolecules by SAS using the Porod-Debye law. Biopolymers, 95(8), 559–571. 10.1002/bip.21638

Sattler, M. (1999). Heteronuclear multidimensional NMR experiments for the structure determination of proteins in solution employing pulsed field gradients. Progress in Nuclear Magnetic Resonance Spectroscopy, 34(2), 93–158. 10.1016/S0079-6565(98)00025-9

Siewering, K., Jain, S., Friedrich, C., Webber-Birungi, M. T., Semchonok, D. A., Binzen, I., Wagner, A., Huntley, S., Kahnt, J., Klingl, A., Boekema, E. J., Søgaard-Andersen, L., & van der Does, C. (2014). Peptidoglycan-binding protein TsaP functions in surface assembly of type IV pili. Proceedings of the National Academy of Sciences of the United States of America, 111(10), E953–61. 10.1073/pnas.1322889111

Skinner, S. P., Fogh, R. H., Boucher, W., Ragan, T. J., Mureddu, L. G., & Vuister, G. W. (2016). CcpNmr AnalysisAssign: a flexible platform for integrated NMR analysis. Journal of Biomolecular NMR, 66(2), 111–124. 10.1007/s10858-016-0060-y

Sørensen, C. S., & Kjaergaard, M. (2019). Effective concentrations enforced by intrinsically disordered linkers are governed by polymer physics. Proceedings of the National Academy of Sciences of the United States of America, 116(46), 23124–23131. 10.1073/pnas.1904813116

Sparling, P. F. (1966). Genetic transformation of Neisseria gonorrhoeae to streptomycin resistance. Journal of Bacteriology, 92(5), 1364–1371. 10.1128/jb.92.5.1364-1371.1966

Stein, D. C., Danaher, R. J., & Cook, T. M. (1991). Characterization of a gyrB mutation responsible for low-level nalidixic acid resistance in Neisseria gonorrhoeae. Antimicrobial Agents and Chemotherapy, 35(4), 622–626. 10.1128/AAC.35.4.622

Stohl, E. A., Chan, Y. A., Hackett, K. T., Kohler, P. L., Dillard, J. P., & Seifert, H. S. (2012). Neisseria gonorrhoeae virulence factor NG1686 is a bifunctional M23B family metallopeptidase that influences resistance to hydrogen peroxide and colony morphology. The Journal of Biological Chemistry, 287(14), 11222–11233. 10.1074/jbc.M111.338830

Stohl, E. A., Dale, E. M., Criss, A. K., & Seifert, H. S. (2013). Neisseria gonorrhoeae metalloprotease NGO1686 is required for full piliation, and piliation is required for resistance to H2O2-and neutrophil-mediated killing. mBio, 4(4). 10.1128/mBio.00399-13

Swanson, J., Kraus, S. J., & Gotschlich, E. C. (1971). Studies on gonococcus infection. I. Pili and zones of adhesion: their relation to gonococcal growth patterns. The Journal of Experimental Medicine, 134(4), 886–906. 10.1084/jem.134.4.886

Swanson, J., Robbins, K., Barrera, O., Corwin, D., Boslego, J., Ciak, J., Blake, M., & Koomey, J. M. (1987). Gonococcal pilin variants in experimental gonorrhea. The Journal of Experimental Medicine, 165(5), 1344–1357. 10.1084/jem.165.5.1344

Szeto, T. H., Dessen, A., & Pelicic, V. (2011). Structure/function analysis of Neisseria meningitidis PilW, a conserved protein that plays multiple roles in type IV pilus biology. Infection and Immunity, 79(8), 3028–3035. 10.1128/IAI.05313-11

Tria, G., Mertens, H. D. T., Kachala, M., & Svergun, D. I. (2015). Advanced ensemble modelling of flexible macromolecules using X-ray solution scattering. IUCrJ, 2(Pt 2), 207–217. 10.1107/S205225251500202X

Trivedi, R., & Nagarajaram, H. A. (2022). Intrinsically Disordered Proteins: An Overview. International Journal of Molecular Sciences, 23(22). 10.3390/ijms232214050

Truebestein, L., & Leonard, T. A. (2016). Coiled-coils: The long and short of it. *BioEssays: News and Reviews in Molecular*, Cellular and Developmental Biology, 38(9), 903–916. 10.1002/bies.201600062

WHO guidelines for the treatment of Neisseria gonorrhoeae. (2016). World Health Organization.

Wilkins, M. R., Gasteiger, E., Bairoch, A., Sanchez, J. C., Williams, K. L., Appel, R. D., & Hochstrasser, D. F. (1999). Protein identification and analysis tools in the ExPASy server. Methods in Molecular Biology (Clifton, N.J.), 112, 531–552. 10.1385/1-59259-584-7:531

Wolfgang, M., van Putten, J. P., Hayes, S. F., & Koomey, M. (1999). The comP locus of Neisseria gonorrhoeae encodes a type IV prepilin that is dispensable for pilus biogenesis but essential for natural transformation. Molecular Microbiology, 31(5), 1345–1357. 10.1046/j.1365-2958.1999.01269.x

