## Supplementary Data for "Structural and functional examination of *Neisseria gonorrhoeae* TfpC reveals a dynamic two-domain architecture required for type IV pilus function"

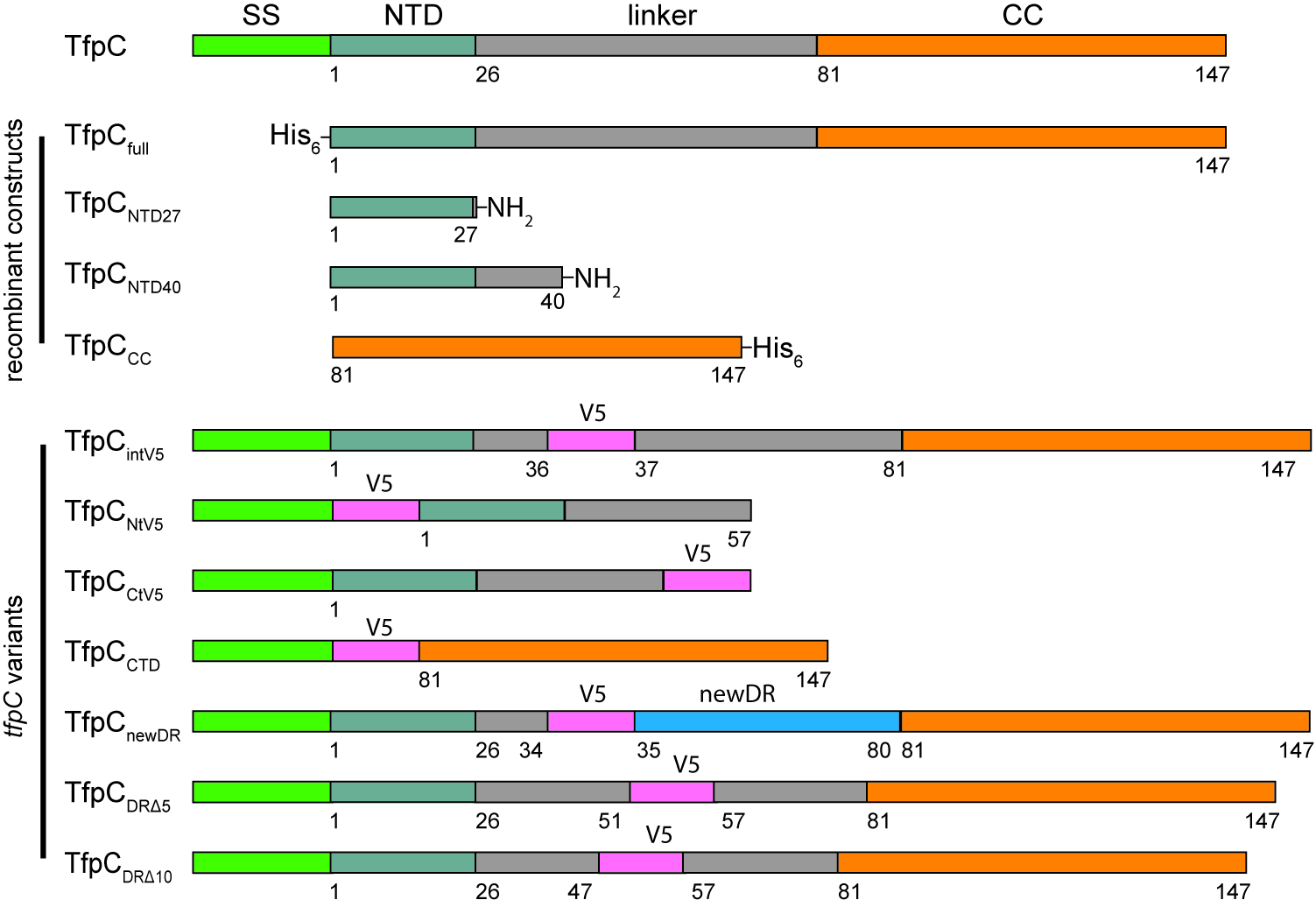

**Supplementary Figure 1: Schematic of Gc TfpC domains and constructs/variants used in this study.** Upper: TfpC is shown with residue numbering based on the mature protein sequence and structural features highlighted. SS: signal sequence; NTD: N-terminal domain; linker: flexible inter-domain linker region; CC: C-terminal coiled-coil domain. Middle: recombinant constructs used in this study with the position of His_6_ affinity tags shown. Peptide modifications are annotated (NH_2_: amidation) along with numbering. Lower: genetic variants used in this study with V5 tag and newDR insertions, and truncations shown, along with numbering.

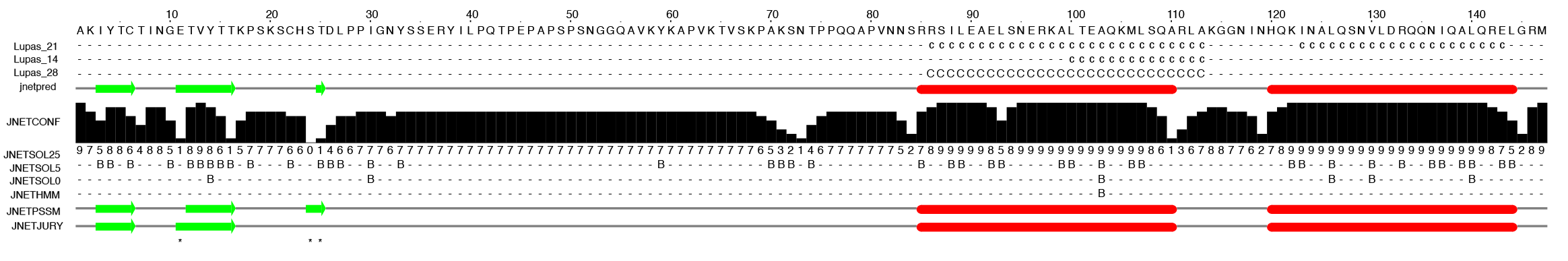

**Supplementary Figure 2. Secondary structure prediction for Ng TfpC.** Jpred4 server (Drozdetskiy et al., 2015) output. Strands are green; helices are red.

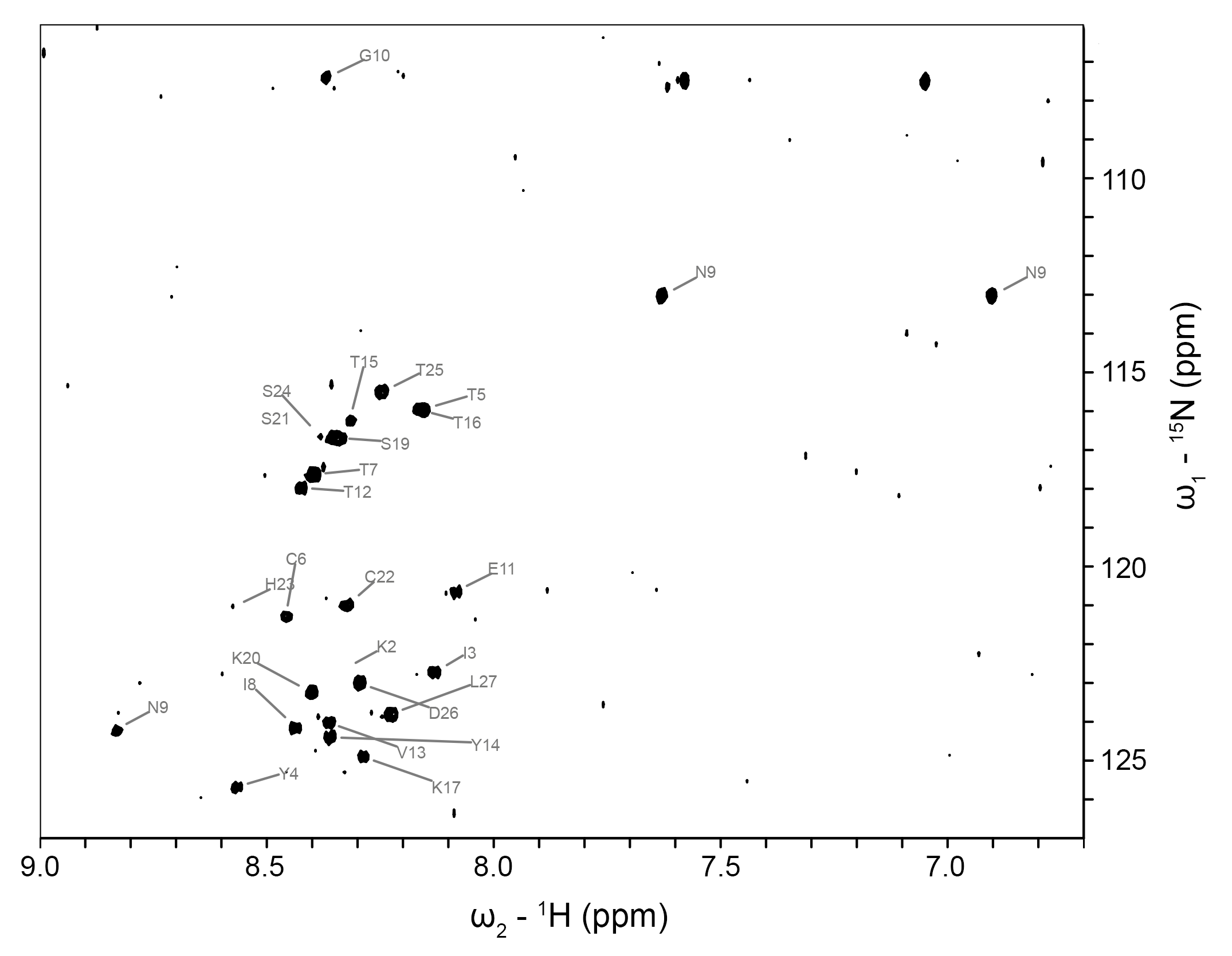

**Supplementary Figure 3: Natural abundance 2D ^1^H^15^N HSQC NMR spectrum of Ng TfpC_NTD27_.** Backbone and amide side chain assignments are annotated.

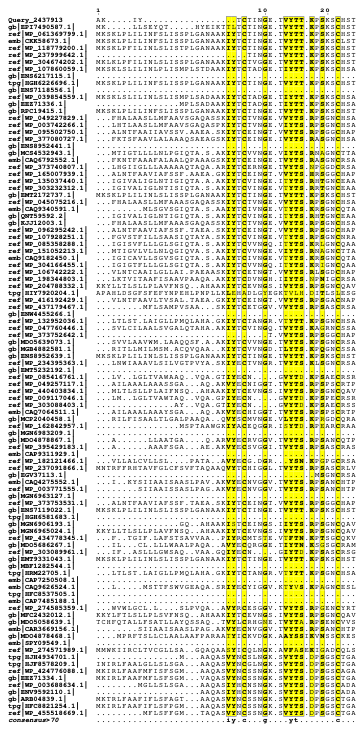

**Supplementary Figure 4. Gc TfpC NTD multiple sequence alignment.** Sequence alignment carried out with NCBI Constraint-Based Multiple Alignment Tool (COBALT) and visualized in the ESPript 3 web server (Gouet et al., 1999). Conserved positions highlighted in yellow.

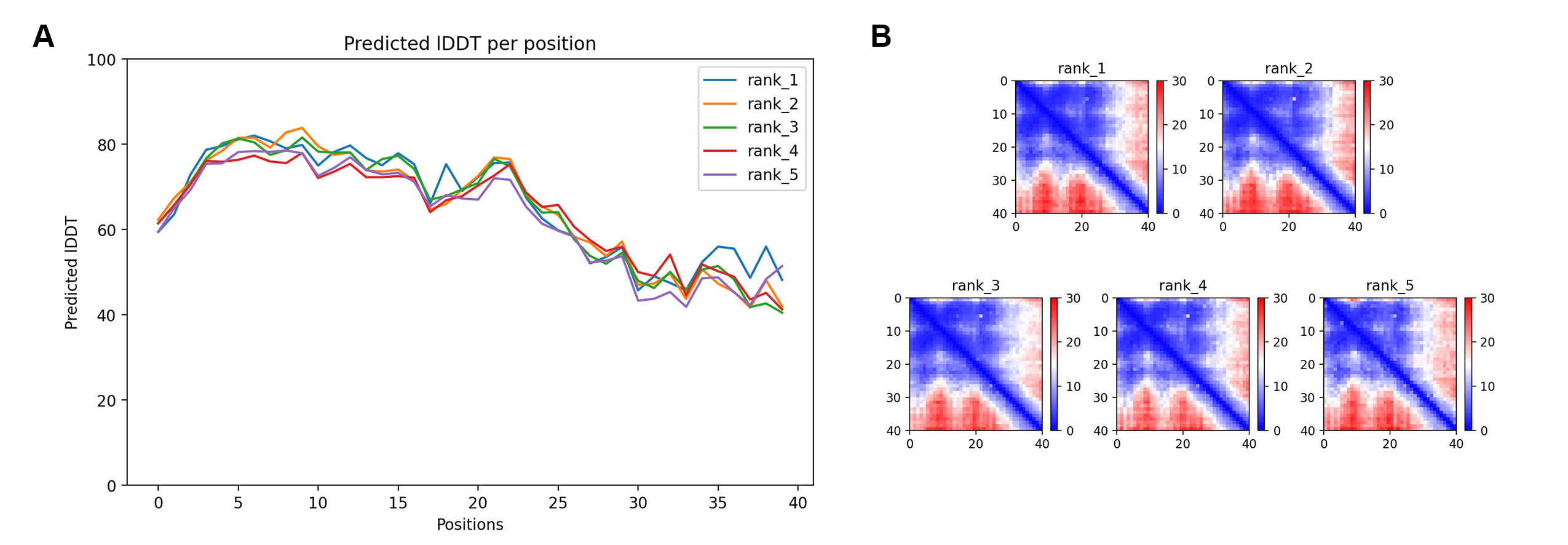

**Supplementary Figure 5: AlphaFold3 modelling of TfpC_NTD40_. (A)** Local Distance Difference Test (lDDT) analysis against residue position for the five top ranking models. High lDDT values (<90) indicate a high accuracy in atomic positions for that residue. Lower values (<70) indicate low confidence atomic positioning, including those observed in dynamic regions. **(B)** Predicted Aligned Error (PAE) analysis of the five top ranking models. Low PAE values throughout the matrix indicates a well folded single domain.

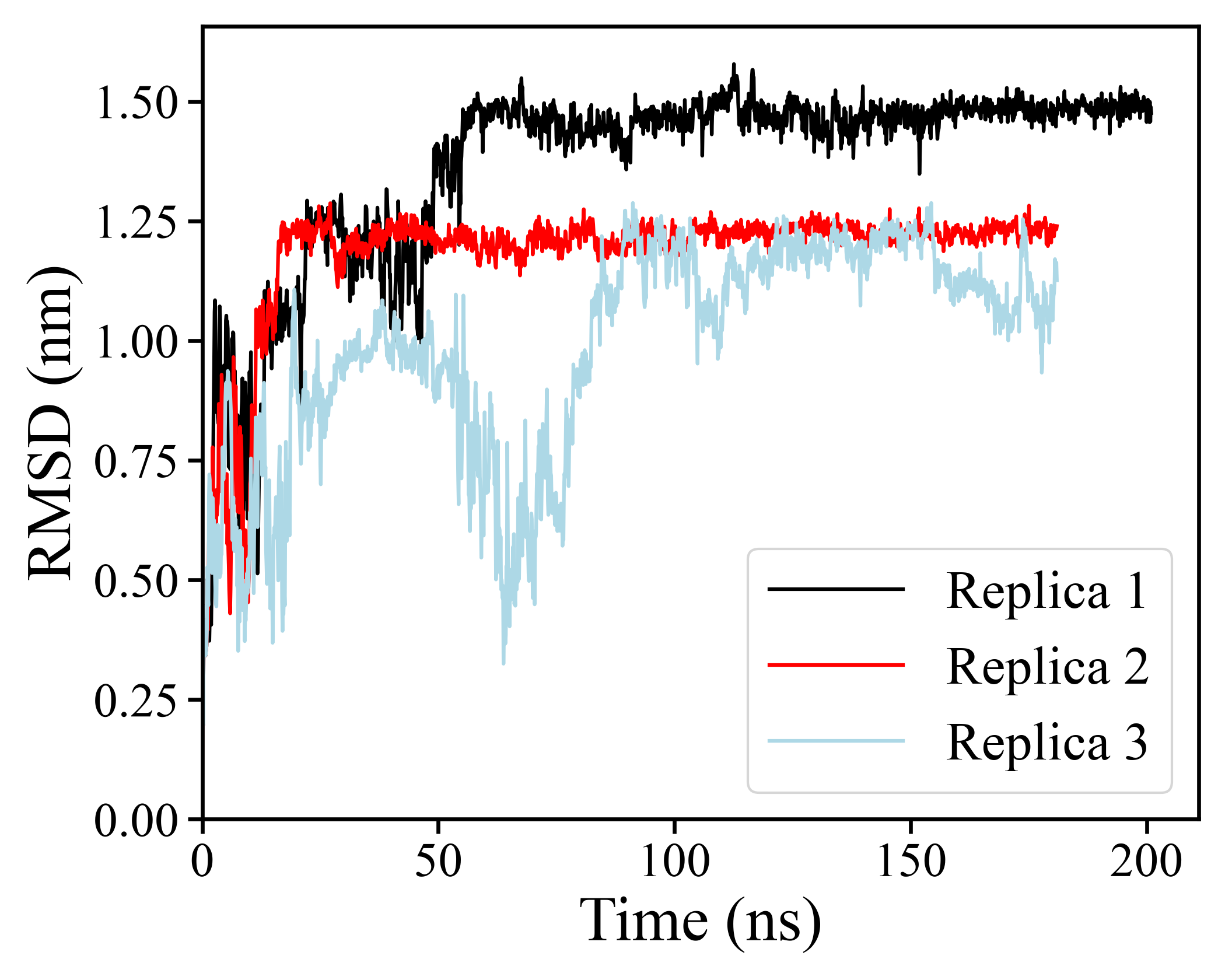

**Supplementary Figure 6. MD simulations of Ng TfpC_NTD40_.** Time evolution of the RMSD of C_a_ atoms from the starting structure before energy minimisation. Simulations were run for 200 ns, followed by two 180 ns replicas.

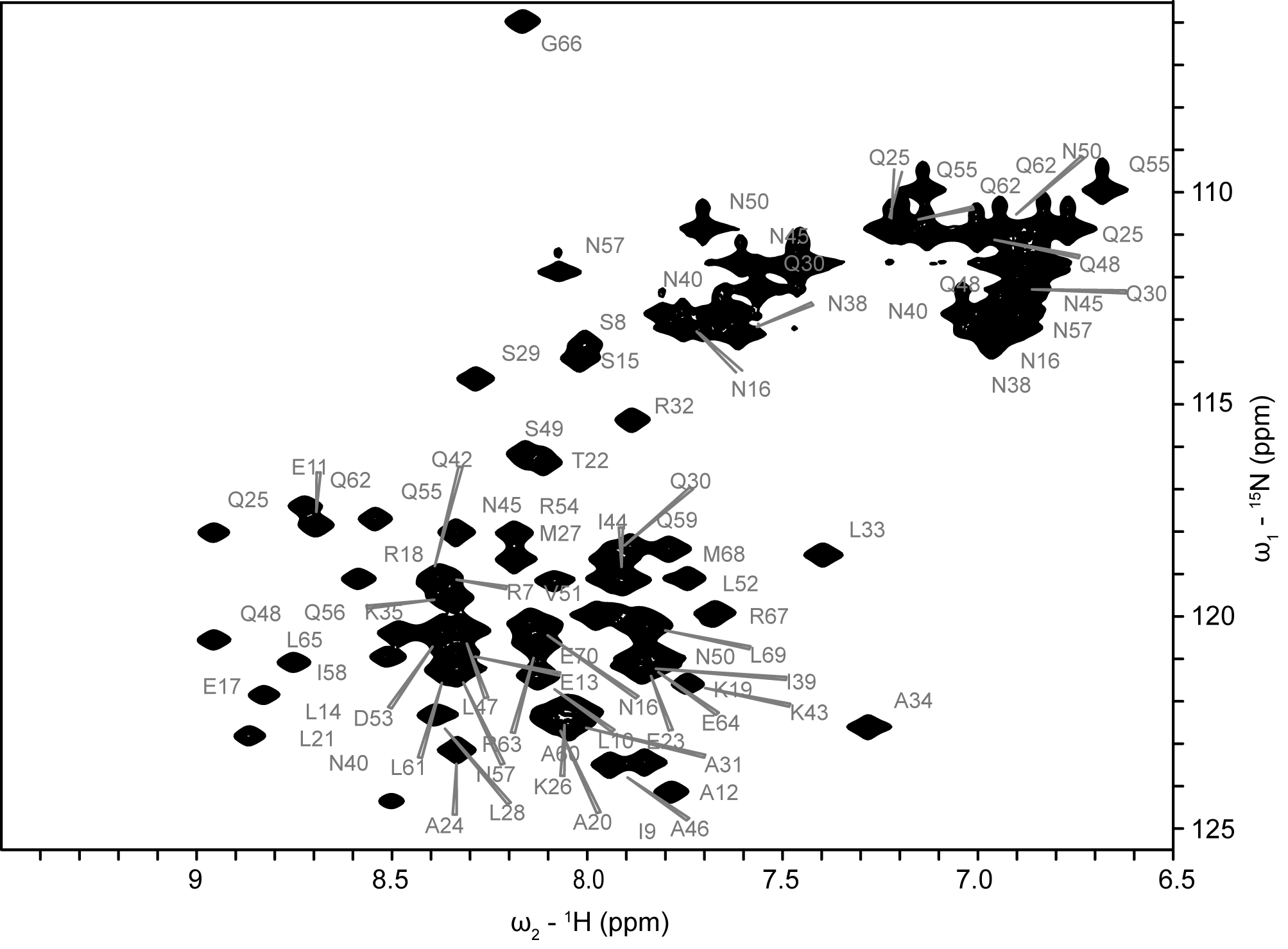

**Supplementary Figure 7: 2D ^1^H^15^N HSQC NMR spectrum of ^15^N labelled Ng TfpC_CC_.** Backbone and amide side chain assignments are annotated.

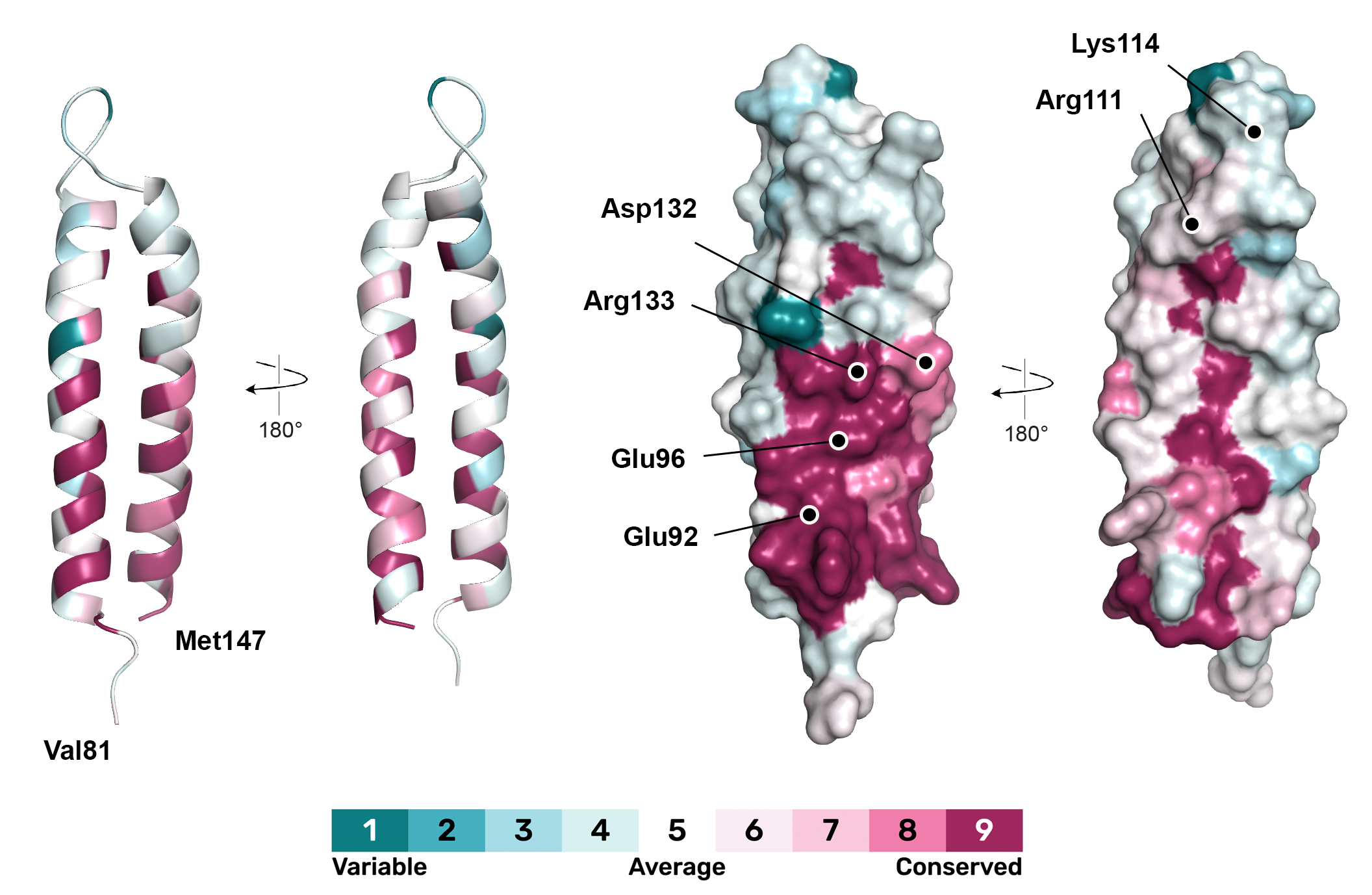

**Supplementary Figure 8: Conservation surface mapping of Ng TfpC_CC_.** Cartoon and surface representation of TfpC_CC_ coloured by conservation, calculate using the ConSurf server (Ben Chorin et al., 2020).

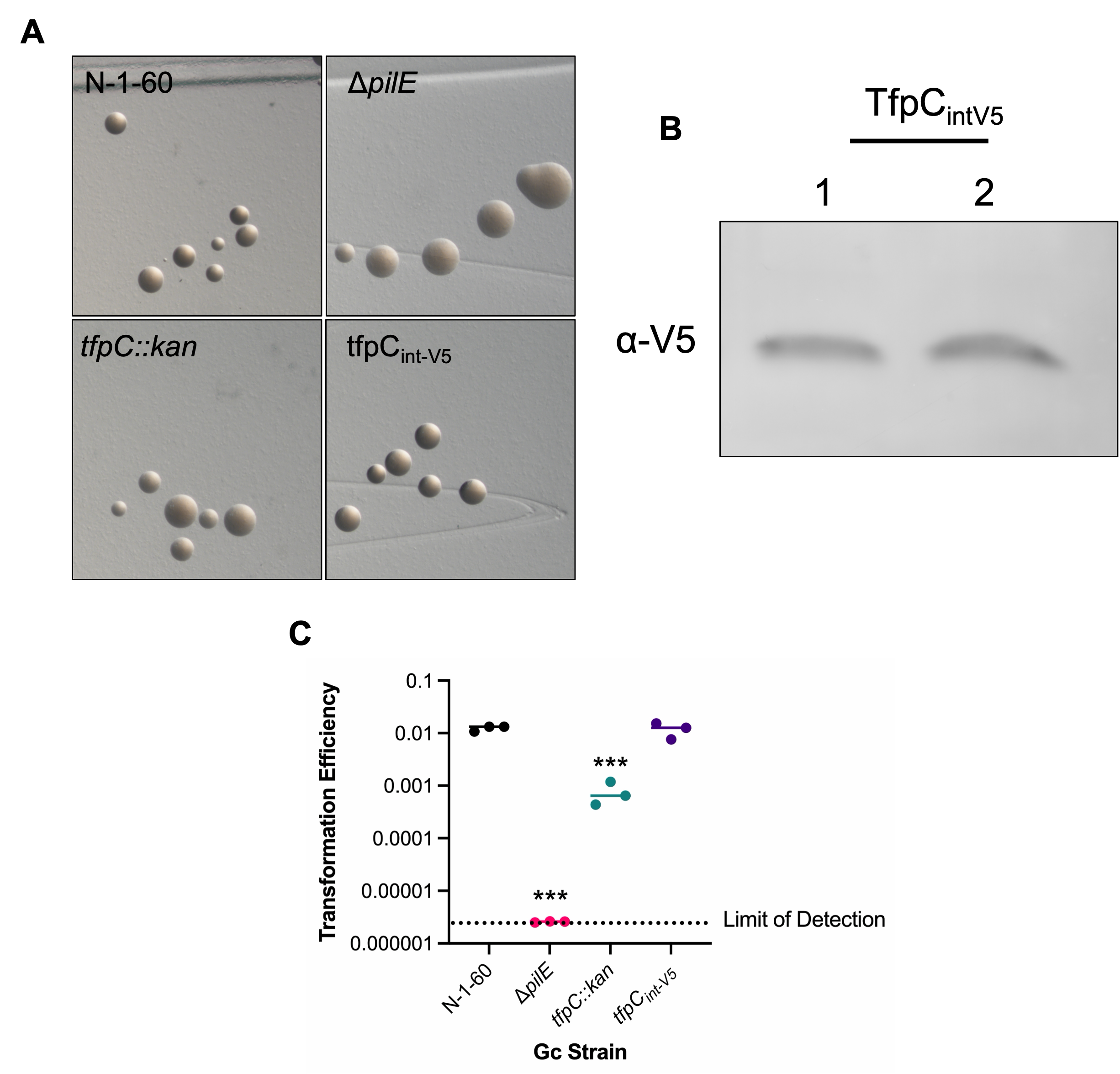

**Supplementary Figure 9: Addition of an internal V5 tag does not alter TfpC protein production or function.** (**A**) Colony morphology assays. (**B**) Western blot of Gc whole cell lysates (40 μg protein load). V5 detects TfpC. (**C**) Transformation efficiency assays were performed using 50 ng of pSY6 plasmid DNA, which confers resistance to nalidixic acid. Cells were treated with DNA for 20 minutes followed by a 10-minute DNase I treatment. Efficiency was calculated by comparing the number of nalidixic acid-resistant cells to the total number of cells plated. Statistical analysis was performed using a one-way ANOVA with Dunnett’s multiple comparisons to the parental strain. ***, p = 0.0002; ***, p = 0.0004.

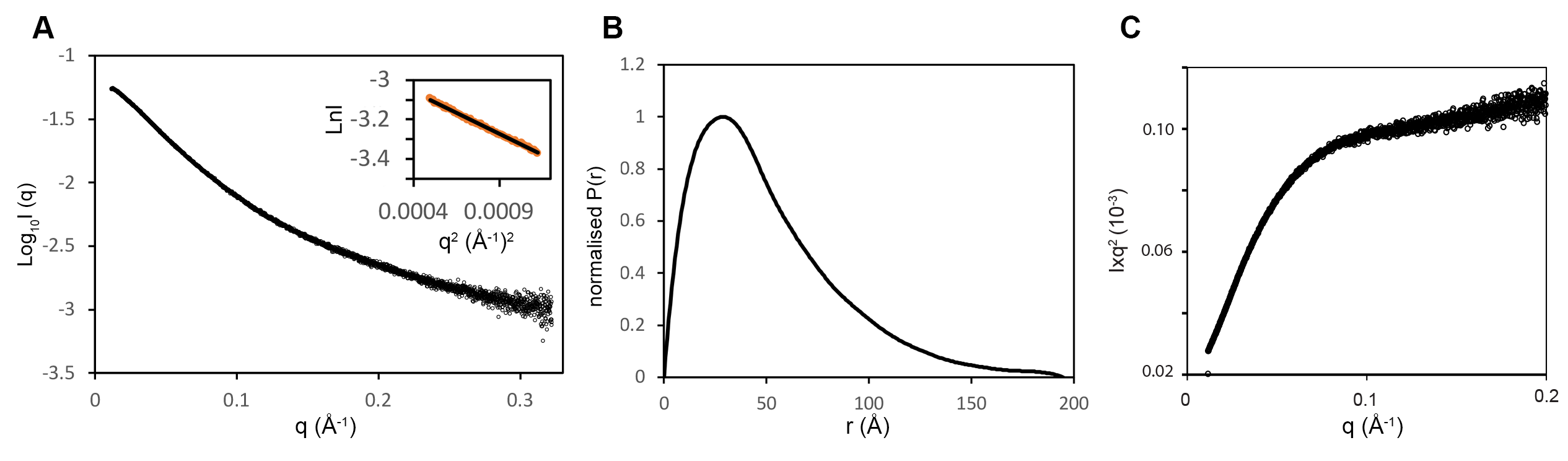

**Supplementary Figure 10: SAXS analysis of Ng TfpC_full_. (A)** Experimental scattering curve of TfpC_full_ (black open circles). Inset: Guinier Region (orange open circles) and linear regression (black line) for R_g_ evaluation. (**B**) Shape distribution [P(r)] function derived from SAXS analysis for TfpC_full_. (**C**) Kratky plot of TfpC_full_.

**Supplementary Table 1 | Summary of NMR Structural Restraints and Statistics**

| **Distance restraints** | |  |
| --- | --- | --- |
|  | Total NOE | 1819 |
|  | Intraresidue | 470 |
|  | Sequential (\|*i* − *j*\|) = 1) | 397 |
|  | Medium range (\|*i* − *j*\|) < 4) | 560 |
|  | Long range (\|*i* − *j*\|) > 5) | 245 |
|  | Ambiguous | 147 |
|  | Dihedral angle restraints | 110 |
|  | *φ* | 55 |
|  | *ψ* | 55 |
|  | Hydrogen bonds restraints | 206 |
| **Structural statistics*^a^*** | | |
|  | **Violations** | |
|  | Distance restraints (NOE) (Å) | 0.083 ± 0.003 |
|  | H-bond Distance restraints (H-bond) (Å) | 0.073 ± 0.002 |
|  | Dihedral angle restraints (°) | 0.0 ± 0.0 |
|  | **Deviation from idealized geometry** | |
|  | Bond length (Å) | 0.008 ± 0.0001 |
|  | Bond angle (°) | 0.90 ± 0.016 |
|  | Impropers (°) | 0.83 ± 0.015 |
|  | **Average pairwise rmsd (Å)** | |
|  | Heavy atoms | 0.50 ± 0.12 |
|  | Backbone atoms | 0.12 ± 0.03 |
| **Structural quality*^b^*** | | |
|  | **Ramachandran statistics** | |
|  | Favored regions (%) | 78.4 |
|  | Allowed regions (%) | 94.6 |
|  | Disallowed regions | 4 residues***^c^*** |

*^a^* Average values and standard deviations over 20 lowest energy conformers with respect to average structure.

*^b^* Percentage of residues in the Ramachandran plot regions determined using an average structure from the 20 lowest energy conformers.

*^c^* All outliers are situated in highly dynamic regions.

**Supplementary Table 2 SAXS data and refinement statistics**

| Beamline | DLS B21 |
| --- | --- |
| Wavelength (Å) | 1.0 |
| q Range (Å^-1^) | 0.12 to 0.34 |
| Structural parameters |  |
| I(0) | 0.055 ± 9.4e-05 |
| R_g_ (nm)  (from Guinier) | 3.56 ± 0.01 |
| R_g_ (nm)  (from P(r)) | 4.36 ± 0.17 |
| D_max_ (nm)  (from P(r)) | 19.5 |

**Supplementary Table 3: SAXS ensemble optimization parameters.**

| R_flex_ (random) (%) | 88.5 ± 0.4 (83.7 ± 0.0) |
| --- | --- |
| R_sig_ | 1.4 ± 0.0 |
| χ^2^ | 2.8 ± 0.1 |
| Cluster 1 (%) | 56 |
| Cluster 2 (%) | 44 |
| Final ensemble <R_g_> (nm) | 38.8 ± 0.4 |
| Final ensemble <D_max_> (nm) | 11.7 ± 0.4 |

**Supplementary Table 4 | Strains**

| Bacterial strain | Serogroup/ genotype | Reference |
| --- | --- | --- |
| BL21 (DE3) | fhuA2 [lon] ompT gal (λ DE3) [dcm] ∆hsdS λ DE3 = λ sBamHIo ∆EcoRI-B int::(lacI::PlacUV5::T7 gene1) i21 ∆nin5 | NEB |
| N-1-60 | FA1090 strain containing a mutation in the guanine quadruplex (G4) upstream of the *pilE* gene preventing antigenic variation | (Hu et al., 2020) |
| Δ*pilE* | N-1-60 strain with an unmarked deletion of the *pilE* gene | (Hu et al., 2020) |
| *tfpC::kan* | N-1-60 strain with an insertional deletion of the *NGO_0783* (*tfpC*) gene. Kan^R^ | (Hu et al., 2020) |
| TfpC_cys🡪ala_ | N-1-60 strain with two single amino acid mutations (aa6 and aa22 of the mature protein) in the *tfpC* gene converting cysteine🡪alanine residues. | This study. |
| TfpC_int-V5_ (“Parental”) | N-1-60 strain with an internal V5 tag. | This study. |
| TfpC_NTD_ | N-1-60 strain expressing the *tfpC* N-terminal domain (aa1-57) with an N or C-terminal V5 tag at the native locus under control of the native promoter. | This study. |
| TfpC_CTD_ | N-1-60 strain expressing the *tfpC* C-terminal domain (aa104-170) with the TfpC signal sequence and an N-terminal V5 tag at the native locus under control of the native promoter. | This study. |
| TfpC_newDR_ | TfpC_int-V5_ strain harboring a mutation in the *tfpC* gene in which the disordered region (aa35-80 of mature protein) has been replaced with the following disordered sequence: QQSKAKAMHRHLTMAMGSDSVQMEAANTNT  HSTANEEDSRAMRANP | This study. |
| TfpC_DRΔ5_ | TfpC_int-V5_ strain containing a five amino acid deletion (aa52-56 of mature protein) within the disordered region. | This study. |
| TfpC_DRΔ10_ | TfpC_int-V5_ strain containing a five amino acid deletion (aa47-56 of mature protein) within the disordered region. | This study. |
| TfpC_E92A/E96A_ | TfpC_int-V5_ strain containing alanine point mutations at residues E92 and E96 of the mature protein. | This study. |
| TfpC_R111A/K114A_ | TfpC_int-V5_ strain containing alanine point mutations at residues R111 and K114 of the mature protein. | This study. |
| TfpC_D132A/R133A_ | TfpC_int-V5_ strain containing alanine point mutations at residues D132 and R133 of the mature protein. | This study. |

**Supplementary Table 5 | Plasmids and primers used in this study**

| Plasmid | Description | Reference |
| --- | --- | --- |
| pTfpCfull | pET28b expression of intact *N. gonorrhoeae* TfpC, residues 1-147 | (Hu et al., 2020) |
| pTfpCcc | pET28b expression of *N. gonorrhoeae* TfpC C-terminal coiled-coil domain, residues 81-147 | This study |
| pTWIST:TfpC_cys🡪ala_ | Plasmid harboring the *tfpC* gene with 1500 bp upstream and downstream of the *tfpC* gene. The *tfpC* gene harbored mutations converting the cysteine residues at aa 6 and aa 22 to alanine residues. | Twist Biosciences |
| pTWIST:TfpC_int-V5_ | Plasmid harboring the *tfpC* gene with 1500 bp upstream and downstream of the *tfpC* gene. The *tfpC* gene harbors an internal V5 tag between residues 37 and 52. | Twist Biosciences |
| pTWIST: TfpC_NTD_ | Plasmid harboring the *tfpC* gene with 1500 bp upstream and downstream of the *tfpC* gene. The *tfpC* gene is truncated to only express the N-terminal domain (aa1-57) with an N or C-terminal V5 tag. | Twist Biosciences |
| pTWIST: TfpC_CTD_ | Plasmid harboring the *tfpC* gene with 1500 bp upstream and downstream of the *tfpC* gene. The *tfpC* gene is truncated to only express the C-terminal domain (aa104-170) with an N or C-terminal V5 tag. | Twist Biosciences |
| pTWIST: TfpC_newDR_ | pTWIST:TfpC_int-V5_ plasmid harboring the *tfpC* gene with 1500 bp upstream and downstream of the *tfpC* gene. This plasmid harbors a mutation in the *tfpC* gene in which the disordered region (aa35-80 of mature protein) has been replaced with the following disordered sequence: QQSKAKAMHRHLTMAMGSDSVQMEAANTNT  HSTANEEDSRAMRANP | Twist Biosciences |
| pTWIST: TfpC_DRΔ5_ | pTWIST:TfpC_int-V5_ plasmid harboring the *tfpC* gene with 1500 bp upstream and downstream of the *tfpC* gene. The *tfpC* gene contains a five amino acid deletion (aa52-56 of mature protein) within the disordered region. | Twist Biosciences |
| pTWIST: TfpC_DRΔ10_ | pTWIST:TfpC_int-V5_ plasmid harboring the *tfpC* gene with 1500 bp upstream and downstream of the *tfpC* gene. The *tfpC* gene contains a five amino acid deletion (aa47-56 of mature protein) within the disordered region. | Twist Biosciences |
| pTWIST: TfpC_E92A/E96A_ | pTWIST:TfpC_int-V5_ plasmid harboring the *tfpC* gene with 1500 bp upstream and downstream of the *tfpC* gene. The *tfpC* gene contains alanine point mutations at residues E92 and E96 of the mature protein. | Twist Biosciences |
| pTWIST: TfpC_R111A/K114A_ | pTWIST:TfpC_int-V5_ plasmid harboring the *tfpC* gene with 1500 bp upstream and downstream of the *tfpC* gene. The *tfpC* gene contains alanine point mutations at residues R111 and K114 of the mature protein. | Twist Biosciences |
| pTWIST: TfpC_D132A/R133A_ | pTWIST:TfpC_int-V5_ plasmid harboring the *tfpC* gene with 1500 bp upstream and downstream of the *tfpC* gene. The *tfpC* gene contains alanine point mutations at residues D132 and R133 of the mature protein. | Twist Biosciences |
| **Primer Name** | **Sequence (5’🡪3’)** |  |
| pilRBS | GGCATTTCCCCTTTCAATTAGGAG | Amplification of the *pilE* gene for sequencing |
| SP3A | CCGGAACGGACGACCCCG |  |
| 135 | TCAGGATCGTGTGCCGATGC | Amplification of the *tfpC* (*NGO_0783*) gene for sequencing |
| 164 | TGTACGTGTAGCCGGTATTGG |  |

**Supplementary Table 6 | Synthetic genes**

| Gene | Description | Sequence (5’ to 3’) |
| --- | --- | --- |
| *tfpCcc* | TfpC C-terminal domain | CCATGGTGAATAATAGCCGTCGTAGCATTCTGGAAGCAGAACTGAGCAATGAACGTAAAGCACTGACCGAAGCACAGAAAATGCTGAGCCAGGCACGTCTGGCAAAAGGTGGTAACATTAATCATCAGAAAATTAACGCCCTGCAGAGCAATGTTCTGGATCGTCAGCAGAATATTCAGGCACTGCAGCGTGAACTGGGTCGTATGCTCGAG |

*Restriction sites are underlined

**References**

Ben Chorin, A., Masrati, G., Kessel, A., Narunsky, A., Sprinzak, J., Lahav, S., Ashkenazy, H., & Ben-Tal, N. (2020). ConSurf-DB: An accessible repository for the evolutionary conservation patterns of the majority of PDB proteins. *Protein Science : A Publication of the Protein Society*, *29*(1), 258–267. https://doi.org/10.1002/pro.3779

Drozdetskiy, A., Cole, C., Procter, J., & Barton, G. J. (2015). JPred4: a protein secondary structure prediction server. *Nucleic Acids Research*, *43*(W1), W389-94. https://doi.org/10.1093/nar/gkv332

Gouet, P., Courcelle, E., Stuart, D. I., & Metoz, F. (1999). ESPript: analysis of multiple sequence alignments in PostScript. *Bioinformatics*, *15*(4), 305–308. https://doi.org/10.1093/bioinformatics/15.4.305

Hu, L. I., Yin, S., Ozer, E. A., Sewell, L., Rehman, S., Garnett, J. A., & Seifert, H. S. (2020). Discovery of a New Neisseria gonorrhoeae Type IV Pilus Assembly Factor, TfpC. *mBio*, *11*(5). https://doi.org/10.1128/mBio.02528-20
